# Apposed networks of interacting TCRs and BCRs exhibiting mosaicked CDR3 sequences made of fixed junctional motifs

**DOI:** 10.1101/2023.05.06.539704

**Authors:** Rizwan Ahmed, Neha Majety, Kevin C. Chan, Adebola Giwa, Joseph Heinemann, Hao Zhang, Joseph Margolick, Rafid Al-Hallaf, Prajita Paul, David R. Bell, Yi Song, Sangyun Lee, Ruhong Zhou, Risa M Wolfe, Thomas Donner, Chunfa Jie, Abdel Rahim A Hamad

**Author notes:** Correspondence to: Abdel Rahim Hamad. and (R.Z). These authors contributed equally (Rizwan Ahmed, Neha Majety).

## Abstract

T and B cells are the two arms of the adaptive immune system that mediate cellular and humoral immunity, respectively, using highly diverse repertoires of antigen receptors. T cells recognize antigens using the T cell receptor (TCR), whereas B cells use the B cell receptor (BCR or surface immunoglobulin). Complementary determining regions (CDR3) of TCRs and BCRs are randomly generated through somatic VDJ recombination and nucleotide deletions and insertions at the V-D and D-J junctions. Contrary to this paradigm, here we describe two networks of millions of TCRβ and IGH clonotypes that are made from only two CDR3 sequences and associated with more 63 diseases. The TCRβ network members bore either the prototypic signature CDR3 sequence (CASSPGTEAFF), its N-terminal VD motif (CASSPGT) recombined with various Jβ segments (CASSPGT-**J**β**x**) or its DJβ motif recombined with various Vβ (**V**β**x**-PGTEAFF). The BCR network members exhibit one signature CDR3 sequence (CARx_1-4_DTAMVYYFYDW) made from an invariant DJH motif (DTAMVYYFDYW) combined with various VH genes. The prototypes of the two networks are apparently teleogically related as they were dually expressed on the rare population of dual expresser (DE) lymphocytes and molecular dynamic simulations show that they were interacting partners. We conclude that members of the two networks represent a core set of evolutionary-conserved primordial antigen receptors that play fundamental roles in host defense and autoimmune diseases.

## INTRODUCTION

T and B cells are the two arms of the adaptive immune system that protect hosts against infections. Each cell type uses highly diverse antigen receptors to recognize various antigens. Consequently, the vast majority of the millions of TCRs and BCRs (surface immunoglobulins, Igs) in different individuals are private (unique to the individual). This is because each antigen receptor is randomly generated by somatic rearrangements of arrays of large numbers of variable (V), diversity (D) and joining (J) genes that are linearly organized in the genome.

Palindromic and random nucleotides added at the V-D (N1 region) and D-J (N2 region) junctions result in further repertoire diversification and paucity of common (public) antigen receptors that are shared by at least by multiple individuals. In concordance, advances in high-throughput technologies have led to uncovering of billions of unique TCR and BCR chains and clonotypes that have been sequenced and deposited in public repositories^1–3^. For example, the iReceptor Gateway Repository alone harbors more than 5 billion sequences generated from hundreds of independent studies and various labs investigating repertoires from infections, vaccinations, autoimmune diseases and healthy individuals^1^. While the vast majority of the published sequences are private clonotypes, public sequences have been identified that are shared among multiple individuals are frequently identified in contexts of various diseases and conditions.

Breaking with these patterns, we here describe two networks of extremely public antigen receptors that were generated from the CDR3 sequences of one public TCRβ and one IGH clonotype, respectively. Besides clonotypes bearing the prototypic signature sequence (CASSPGTEAFF), other members of the TCRβ network bear one of two overlapping sequences of it. In the dominant set, the VβD-motif (CASSPGT-) was recombined with various Jβs, whereas in the minor set, the –DJβ motif (-PGTEAFF) was recombined with different Vβ genes, resulting in signature CDR3 sequences: CASSPGT-Jβx and Vβx-PGTEAFF (x any gene), respectively. The BCR network is made of the -DJH motif (-DTAMVYYFDYW) of the prototypic clonotype (CARDQEDTAMVYYFDYW) recombined with various VH genes, resulting in the signature sequence CARx_1-4_DTAMVYYFYDW (where x is any amino acid). The networks were identified by serendipity while investigating TCRβ repertoires of T cells captured by a public IgM monoclonal antibody (referred to as x-mAb) produced by dual expresser (DE) lymphocytes^4^. Subsequent in-depth analysis of more than 6-billion publicly available sequences led to the identification of the TCRβ network. On the other hand, the IGH chain of the x-mAb turned out to be the prototype of the BCR network. Analysis of these networks could lead to new hypotheses and mechanistic studies to accelerate understanding autoimmune and immune response repertoires of T and B cells.

## RESULTS

### Capturing phenotypically distinct set of T cells using a self-reactive monoclonal antibody

We have recently identified a lymphocyte that coexpresses TCR and BCR and referred to as dual expressers (DEs)^4^. The antibody produced by DEs in patients with type 1 diabetes (T1D) is self-reactive and of the IgM isotype (called the x-mAb)^4^. In our efforts to characterize specificity and pathogenic activity of the x-mAb, we investigated its interacting T cell partners. The x-mAb-reactive T cells were identified by preincubating PBMCs from T1D or healthy controls (HC) subjects (**Supplementary Table 1**) with purified x-mAb followed by staining with fluorochrome-conjugated anti-IgM secondary antibody, anti-TCR, and anti-CD4 or CD8 specific antibodies. FACS acquired data were analyzed using the FlowJo software (**Extended Data 1**). X-mAb-reactive T cells were detected in both T1D and HC subjects (**Fig. 1a** and **Extended Data 2a**). x-mAb-reactive T cells (for brevity referred to as Tx cells in figures and legends) were comprised of both of CD4 and CD8 T cells whose frequencies as percentages of total CD4 or CD8 T cells were significantly higher in T1D than in HC subjects. Interaction of x-mAb was not due to their expression of the Fcα/μR as its blockade using an increasing dose of human serum IgM did not alter binding of directly labeled, AF-488-x-mAb, to T cells (**Extended Data 3**). The results are consistent with published results showing the Fc mu receptor (Fcα/μR) is expressed on the majority of B lymphocytes and macrophages, but not on granulocytes and T cells^5, 6^. Because of the paucity of x-mAb-reactive T cells in HCs, subsequent experiments were limited to x-mAb-reactive T cells in T1D patients, unless stated otherwise. X-mAb-reactive CD4 and CD8 T cells overwhelmingly expressed the CD69 activation maker (**Fig. 1b** and **Extended Data 2b**), and a significant fraction coexpressed CD25 and CXCR3 molecules as compared to Tcon cells (**Fig. 1c** and **Extended Data 2c**). x-mAb-reactive CD4 and CD8 T cells, similar to their Tcon counterparts, however, were comprised of naïve and antigen-experienced (effector memory, TEMRA, and central memory) subsets (**Fig. 1d** **and Extended Data 2d**). The only exception is that of central memory (Tcm) cells, which comprised a significantly higher fraction of x-mAb-reactive T cells relative to Tcon cells especially among the CD8 T cells (**Fig. 1d** **right graph** and **Extended Data 2d**).

**Fig. 1.**
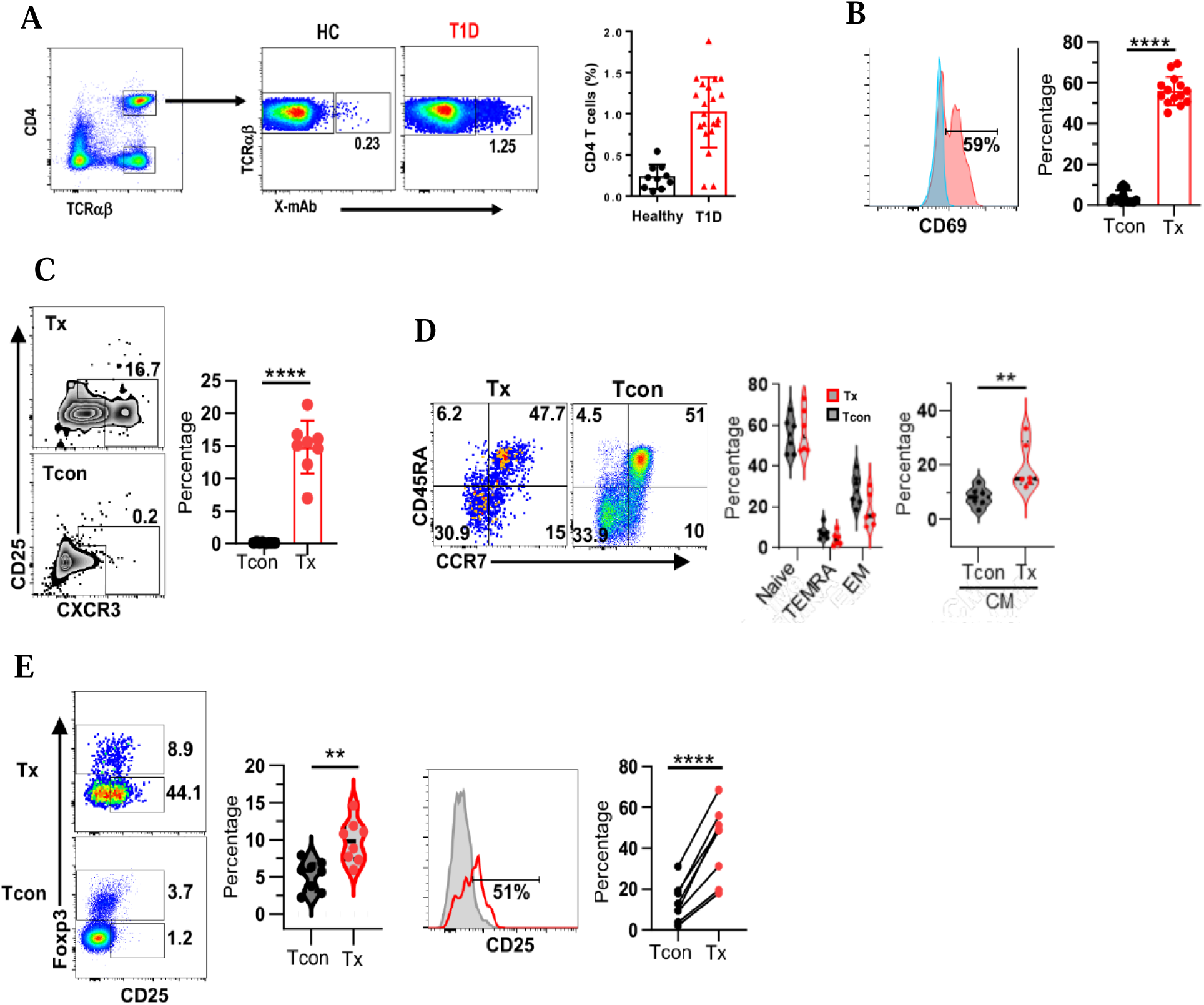
Phenotypic features of x-mAb-reactive CD4 T cells. (a) Representative dot plots show staining of a subset of CD4 T cells by the x-mAb (Tx cells) in HC and T1D subjects. Numbers indicate percentages. **Graph** shows cumulative data (Mean ± SEM; T1D (n=21) and HC (n=10). **P<0.01 by two samples independent t test. (**b**) Histogram overlay shows frequency of CD69^+^ among CD4 Tx (red) and Tcon (blue) cells. Numbers indicate percentage in Tx cells. **Graph** shows cumulative data (n=14 subjects). ****P<0.0001 by paired t test. (**c**) Representative dot plots show expression CD25 and CXCR3 by CD4 Tx and Tcon cells. Numbers indicate percentages. **Graph** shows cumulative data (Mean ± SEM; n=8. ****P<0.0001 by paired t test. (**d**) Representative dot plots showing expression of CD45RA and CCR7 by CD4 Tx and Tcon cells. **Left**, violin graph shows cumulative data of naïve, TEMRA, and effector memory subsets among Tx and Tcon subsets. **Right**, violin graph shows cumulative data (Mean ± sem; n=6) for T_CM_ central memory subset among CD4 Tx and Tcon subsets. **P<0.01 by student’s paired t test. (**e**) Representative dot plot shows Treg cells (Foxp3+CD25+) among gated CD4 Tx and Tcon cells. Numbers indicate percentages. Violin plots show cumulative data Treg cells (%) in each subset. Overlay showing expression of CD25 by gated Tcon (grey) versus Tx (red) subsets. Line graph shows percentage of CD25^hi^ in each subset (Mean ± sem; n=8). **P<0.01, ****P<0.0001 by paired t test.

Consistent with their functional heterogeneity, we detected no significant differences in the frequency of cells expressing key proinflammatory (TNFα, IFN-γ, IL-17) and anti-inflammatory (IL-10) cytokines between x-mAb-reactive and Tcon cells. Albeit, frequency of IL-2 expressing was significantly higher among CD4 Tcon cells than in their x-mAb-reactive counterparts (**Extended Data 4**). Foxp3^+^ CD4 Treg cells were present among Tcon cells, but constituted a significantly larger percentage of x-mAb-reactive T cells, particularly of CD25^hi^ Foxp3^+^ Treg cells (**Fig. 1e**). The results show that the x-mAb interacts with a subpopulation of CD4 and CD8 T cells with distinctive phenotypic features as compared to autologous CD4 and CD8 Tcon cells.

### X-mAb-reactive T cells are anchored by an extremely public TCRβ clonotype

Next, we sorted and determined whether the TCRβ repertoire of x-mAb-reactive T cells is distinct from that of non-reactive autologous Tcon cells using the immunoSEQ assay (**Extended Data 5**). We found that TCRαβ repertoires of sorted x-mAb-reactive T cells across donors to be polyclonal with diverse Vβ, D and Jβ gene usage, a pattern similar to that of Tcon cells (**Supplementary Table 2**). The results are consistent with accumulating evidence of polyclonality of autoreactive T cells. We detected, however, 19 public (shared) clonotypes that exhibited intriguing differences in usage by the two subpopulations (**Fig. 2a**). Specifically, 13 out of the 19 public clonotypes were exclusively shared by Tcon cells whereas 2 other clonotypes were exclusively shared by x-mAb-reactive T cells across donors. On the other hand, 4 clonotypes were commonly shared by Tcon and Tx cells across donors. One of the shared clonotypes (CDR3: CASSYPGTEAFF) stood out because it was enriched 2- to 250-fold in x-mAb-reactive relative to Tcon cells (**Fig. 2b**). To assess the public prevalence and relevance of this clonotype, we investigated its usage in three public database repositories that complement each other: (i) The iReceptor Gateway repository which harbors over 5 billion TCR and BCR sequences^1^. Because of enormous size of deposited data in the iReceptor, its searching engine, however, does not allow identification of related sequences. (ii) The TCR database (TCRdb) repository^2^ whose relatively smaller size (∼277 million TCRβs) allows its searching engine to identify not only identical, but also related clonotypes with up to two amino acid mismatches and it also links clonotypes to specific disease conditions. (iii) The network of Pancreatic Organ Donors with Diabetes (nPOD), which is a T1D-focused repository that harbors TCR and BCR sequences from various organs including pancreatic lymph nodes (PLNs), the site of diabetogenic T cell priming, as well as from disease-irrelevant organs (e.g. irrelevant lymph nodes (iLNs); spleens; PBMCs) of T1D and control subjects^7^.

**Fig. 2.**
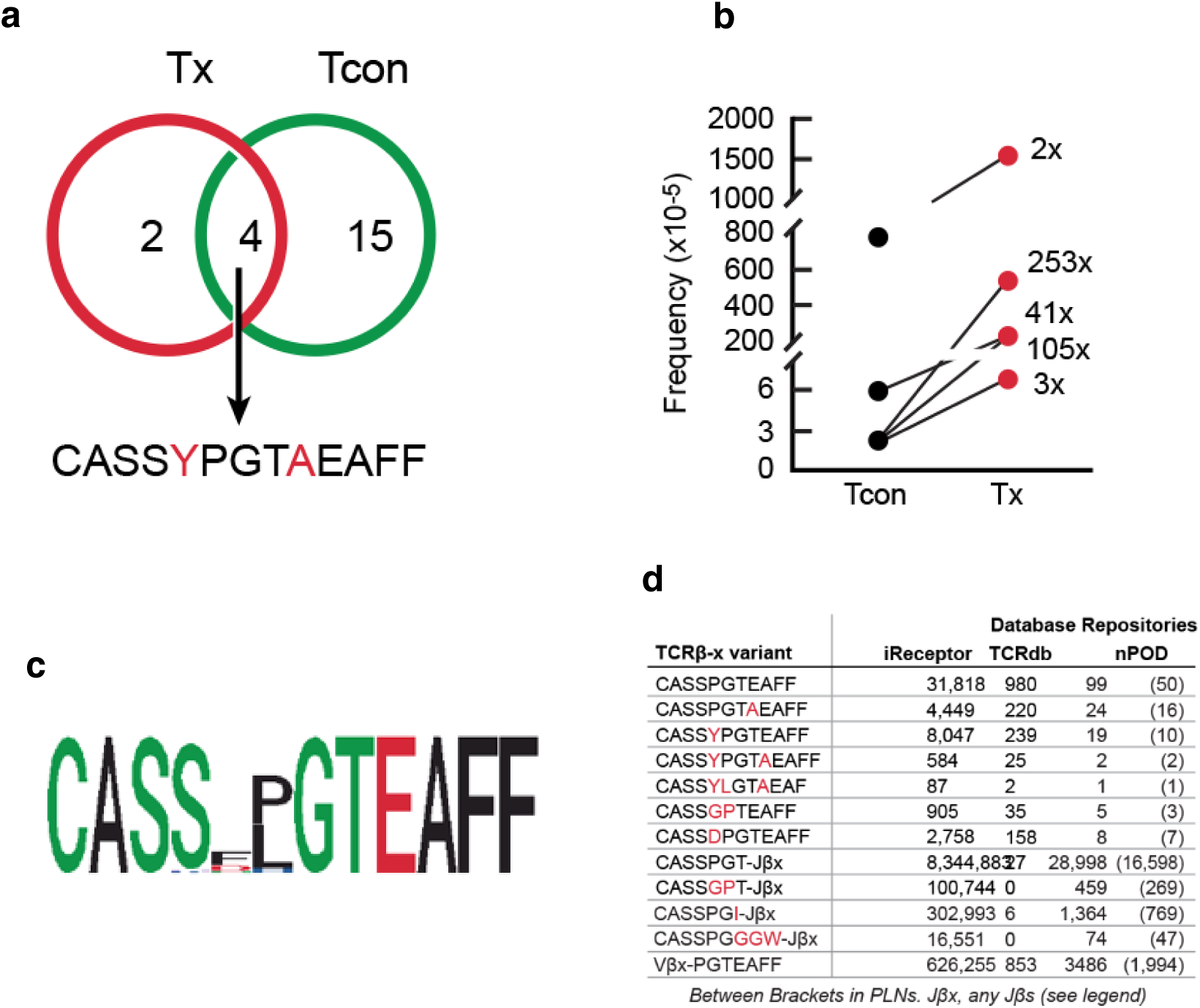
X-mAb-captured public TCRβ clonotype uncovered an elaborate network of highly related TCRβ clonotypes. (**a-b**) Venn diagram depicts numbers of public TCRβ clonotypes that are shared or unique to the Tx or Tcon subpopulations in five T1D-subjects (**a**). Arrow points to the most x-mAb enriched CDR3 amino-acid sequence of the TCRβ-clonotype (red-typed letters indicate variable amino acids). Graph shows the frequency of the clonotype among autologous Tx (red) and Tcon (black) cells per subject (**b**). Enrichment (fold change) of the clonotype in Tx relative to Tcon cells per subject is next to lines. (**c**) Logo depicts the consensus CDR3 sequence of the alignment for the retrieved TCRβ-x family members. (**d**) Table shows numbers and CDR3 sequences of the family prototypic-clonotype (**CASSPGTEAFF**) and indicated family members that were retrieved from the iReceptor, TCRdb and nPOD repositories. Table also shows total numbers of clonotypes bearing the other two signature (**CASSPGT-J**β**x**) or DJβ (**V**β**x-PGTEAFF**) sequences. Letter x in Jβxs and Vβx refers to any Jβ and Vβ gene in the genome, respectively (**see Supplementar**^1^**y**^8^ **Tables**).

Our database search revealed that this x-mAb-interactive clonotype as a ubiquitously used public clonotype that has numerous variants that differed mostly by a single residue yielding a canonical (CDR3: CASSxPGTxEAFF) sequence with x being any residue (**Fig. 2c**). Variable residues were notably inserted either before and/or after the PGT tripeptide, which is conserved in most clonotypes. In addition, the most abundant clonotype (CDR3: CASSPGTEAFF) has no insertions flanking the PGT tripeptide and considered the prototype (hereafter referred to as the TCRβ-x) of the group. The TCRβ-x clonotype had 31,818 occurrences in the iReceptor Gateway database (3,426 repertoires, 1,913 subjects, 12 tissues, 26 diagnoses, 23 research labs and 30 studies), 980 occurrences in the TCRdb database, and 99 occurrences in the T1D-focused nPOD repository (**Fig. 2d** and **Supplementary Tables 3-4**). Several variants of the TCRβ-x clonotype are also highly prevalent public antigen receptors with hundreds of thousands of occurrences (**Fig. 2d**). The TCRβ-x prototype and variants were used by both CD4 and CD8 T cells, indicating lack of MHC restriction and wide usage. In line with this notion, the 980 TCRβ-x clonotypes identified in the TCRdb (**Extended Data 6a** and **Supplementary Table 4**) were linked to 63 diseases and conditions (autoimmune diseases led by T1D, cancers and infections). Moreover, we identified the TCRβ-x clonotype among high-confidence SARS-CoV-2-specific TCRs^8^ and found that it was associated with recognition of seven COVID-19 epitopes (**Extended Data 6b**). Among nPOD subjects (**Supplementary Table 5**), the TCRβ-x clonotype was used by CD4, Treg and CD8 T cells isolated from PLNs, spleens and iLNs of both T1D (16 subjects, 23 organs, 53 sequences) and control (7 subjects, 13 organs, 28 sequences) subjects. In both T1D and control subjects, approximately half of the TCRβ-x clonotypes were from PLNs and 83% of CD4 T cells isolated from PLNs of controls were of the Treg lineage compared to the 46% of CD4 T cells in isolated PLNs of T1D subjects being of Treg lineage (**Extended Data 7**). Thus, the TCRβ clonotype most enriched by the x-mAb is associated with various diseases and used by both CD4 and CD8 T cells, including those in PLNs of T1D subjects.

### Identification of a unique TCRβ network built from overlapping motifs of the VDJ sequence of the TCRβ-x prototype

While identical at the nucleotide level in various individuals, the CDR3 sequence (CASSPGTEAFF) of the TCRβ-x clonotype is encoded by any of 20 Vβ genes (**Fig. 3a**) recombined to the same DJβ motif encoded by the TRBD1*01 and TRBJ1-01*01 genes (<2-10% used TRBD2). In each clonotype, the Vβ gene contributed to the nucleotide sequence encoding the PGT tripeptide junction. N1- and P-nucleotide additions and 5‣ and 3‣ trimmings of the TRBD1*01 gene kept the CDR3 sequence in frame in different clonotypes. Examples highlighting how different Vβs were used to generate the same CDR3 sequence and the PGT tripeptides are shown (**Fig. 3a**). Another highly related and ubiquitous CDR3 sequence, CASSGPTEAFF (note the reversed G and P positions), is generated by various Vβ gene recombinations to the TRBD1*01-TRBJ1-01*01 motif (**Supplementary Tables 6a-c**). Thus, the CDR3 sequence of the TCRβ-x clonotype is generated by different combinations using different Vβ genes recombined to the same DJβ motif.

**Fig. 3.**
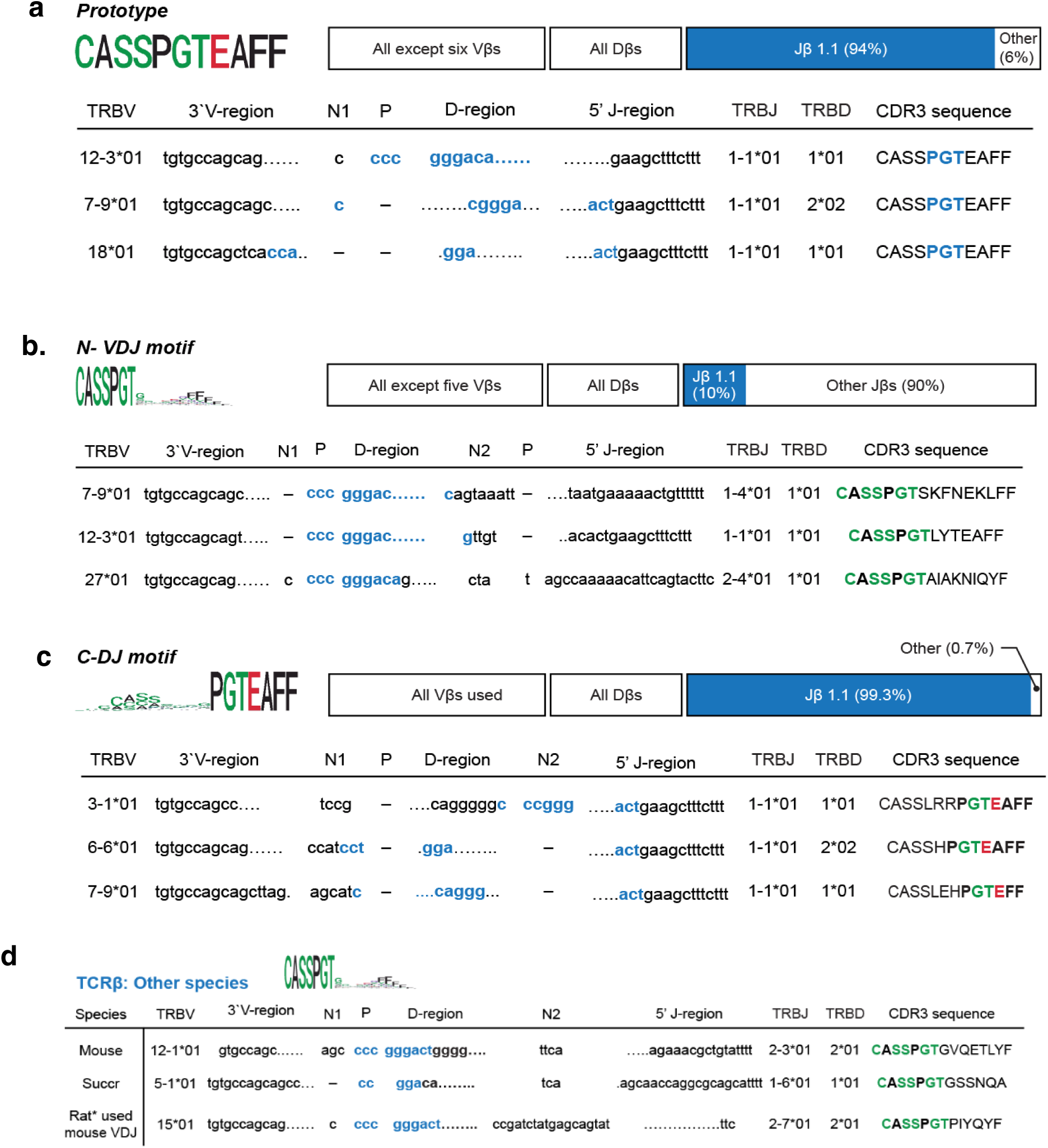
Examples of how overlapping motifs of the VDJ sequence of the TCRβ prototype are used to generate the network. (**a**) Logo and Diagram show usage of various Vβ and D genes with mostly Jβ1 gene to generate the prototypic signature sequence (**CASSPGTEAFF**). (**b-c**) Logos and diagrams are representative examples depicting usage of various Jβs with the VβD (**CASSPGT**) motif to generate clonotypes bearing the **CASSPGT-J**β**x** signature sequence (**b**) or various Vβs with the DJβ (**PGTEAFF**) to generate the clonotypes bearing the **V**β**x-PGTEAFF** signature sequence. (**c**) Amino acids comprising the NDJ junction in each TCRβ group in humans are in blue. (**d**) Table shows examples of V, D and J genes used for generating CDR3-sequences of hybrid TCRβ clonotypes using the VD-motif in mice, pigs (Suc Scrofa) and rat.

Furthermore, the CDR3 amino acid sequence of the prototypic TCRβ-x clonotype is broken down into two overlapping motifs, a VD-(CASSPGT-) and a –DJ motifs (-PGTEAFF), that are systemically used to generate partly identical sequences. The CASSPGT-motif was rearranged to various Jβs, generating clonotypes defined by a CASSPGT-Jβx (x represents any Jβ) CDR3 signature sequence (**Fig. 3b**). Conversely, the -PGTEAFF fragment was rearranged to various Vβ genes resulting in a set of clonotypes with a Vβx-PGTEAFF (x any of most Vβ genes) CDR3 signature sequence (**Fig. 3c**). In total, we identified 8,344,833 CASSPGT-Jβx clonotypes (6698 repertoires, 2930 subjects, 4572 samples, 48 diagnoses, 31 tissues) from the iReceptor repository alone (due to sheer size sequences can be viewed at the repository site). Another set of 6,196 CASSPGT-Jβx sequences were retrieved from the nPOD database (**Supplementary Table 7**). TCRβ clonotypes exhibiting the CASSPGT-Jβx CDR3 sequence were detected in mice, pigs, and Lewis rats using Blastp search (**Fig. 3d**), indicating versatile and evolutionary conserved usage. On the other hand, we detected 626,255 Vβx-PGTEAFF hybrid clonotypes in the iReceptor repository (**Supplementary Table 8a**) and 3,486 in the nPOD database (**Supplementary Table 8b**). Thus, the CDR3 sequence of the TCRβ-x clonotype is made of two overlapping VβD and DJβ motifs that are used independently yet systemically to generate a network of an extremely large number of hybrid clonotypes sharing the N- or C-terminal domains including the PTG coding joint.

### The VD and DJ motifs of TCRβ-x are used by IGH clonotypes and proteins encoded by single genes

An additional three TCRβ clonotypes (CASSGPT-Jβx, CASSPGI-Jβx, and CASSPGGGW-Jβx) with hundreds of thousands of occurrences were identified (**Supplementary Table 9a-b**; **Supplementary Table 10; Supplementary Table 11).** Intriguingly, the same VD-motifs were used by BCR heavy chains, generating IGHV clonotypes exhibiting CDR3 sequences (CASSPGT-JHx, CASSGPT-JHx, CASSPGI-JHx, and CASSPGGGW-JHx) corresponding to those of the TCRβ clonotypes (**Fig. 4a** and **Supplementary Tables 12a-d**). These observations show that the same amino acids representing VD motifs of the TCRβ-x clonotypes are used in the CDR3 sequences of not only TCRs but some BCRs. While unprecedented in humans, to the best of our knowledge, the TCRµ chain of marsupials is considered a chimera between the IGH and TCR loci that uses prejoined VD and VDJ segments^9^. Trans-rearrangements of IGHV segments with TCR gene loci were also reported in the nurse shark^10^, suggesting structural similarities of the TCRβ-x sequence to the ancient receptor systems of elasmobranch. It is therefore interesting that the VD-(CASSPGT) and -DJ (PGTEAFF) motifs are used by enzymes and transporters in bacteria and humans (**Fig. 4b-c** and **Supplementary Table 13**). Thus, by analysis of the CDR3 sequence of the TCRβ most enriched by x-mAb, we demonstrate existence and wide usage of ancient VD and DJ motifs by public TCRβs and IGHVs.

**Fig. 4.**
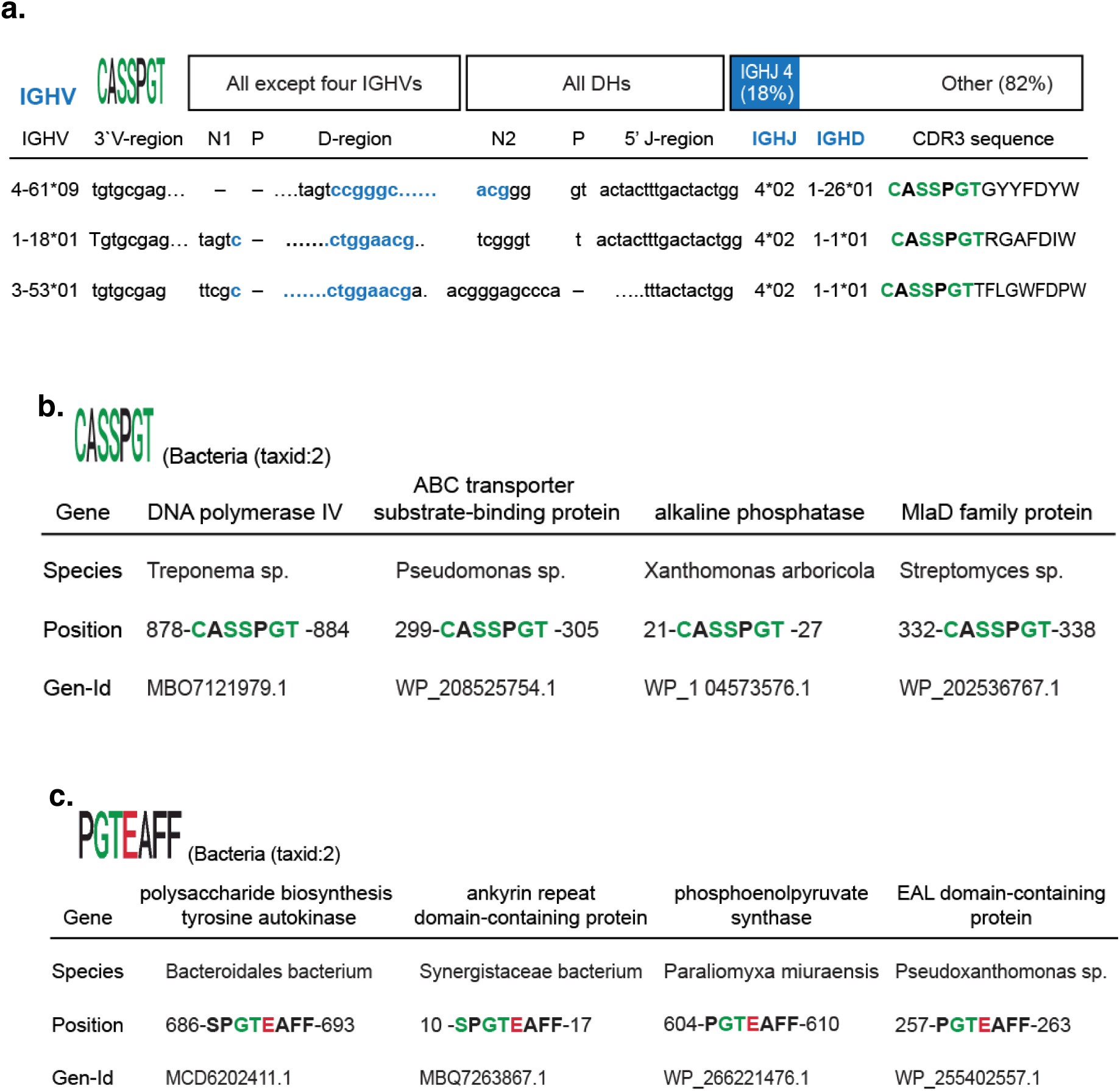
The VβD and DJβ motifs of TCRβ-x are used by IGH clonotypes and one gene-encoded bacterial proteins. (**a**) Table shows examples of VH genes used to generate hybrid IGHV clonotypes bearing the VD-motif. Blue letters represent amino-acid residues comprising VDJ junctions in different clonotypes. (**b**) Table shows usage of the **CASSPGT** sequence motif by bacterial enzymes in indicated species. (**c**) Table shows usage of the **PGTEAFF** sequence-motif by bacterial enzymes in indicated species.

### Association of public TCRα clonotypes by members of the TCRβ-x network

A TCRβ clonotype (CDR3: CASSYLGTAEAFF) that we cloned from DE cells is closely related to the TCRβ-x prototype^4^. It is paired on DEs with a TCRα clonotype that was originally referred to as the TCRα-x (CDR3: CAASASGGGGSNYKLTF). High-throughput analysis showed the TCRα-x clonotype was highly enriched (up to >2000-fold) in DEs relative to autologous Tcon cells (**Fig. 5a-b** and **Supplementary Table 14a**). Furthermore, analysis of TCRα repertoires of x-mAb-reactive and non-reactive Tcon cells from a T1D subject identified the TCRα-x at the top TCRα clonotype in x-mAb-reactive T cells compared to being #72952 in autologous Tcon cells, indicating selective enrichment by the x-mAb (**Supplementary Table 14b**). We identified 72 occurrences of the TCRα-x and closely related clonotypes that differed by missing one glycine residue at the N/Jα53 junction in the iReceptor database (CAASAS*GGGSNYKLTF). An additional 1,125 clonotypes (71 repertoires, 4 studies, 23 subjects, 71 samples, 5 diagnoses, 4 tissues) shared the coding sequence (SGGGGSNYKLTF) generated by N region additions and the J53α gene (**Fig. 5c** and **Supplementary Table 15**). The TCRα-x clonotype is closely related to two insulin-reactive murine TCRα clonotypes (CDR3: Vα5-4*01-CAASASGSGGSNYKLTF-Jα53; CDR3: Vα5-4*01-CAASASGGSNYKLTF-Jα53) that were cloned from T cells isolated from pancreatic infiltrates of NOD mice; and were considered drivers of the diabetogenic process based on their ability to induce anti-insulin autoimmunity in transgenic and retrogenic NOD mice^11–13^.

**Fig. 5.**
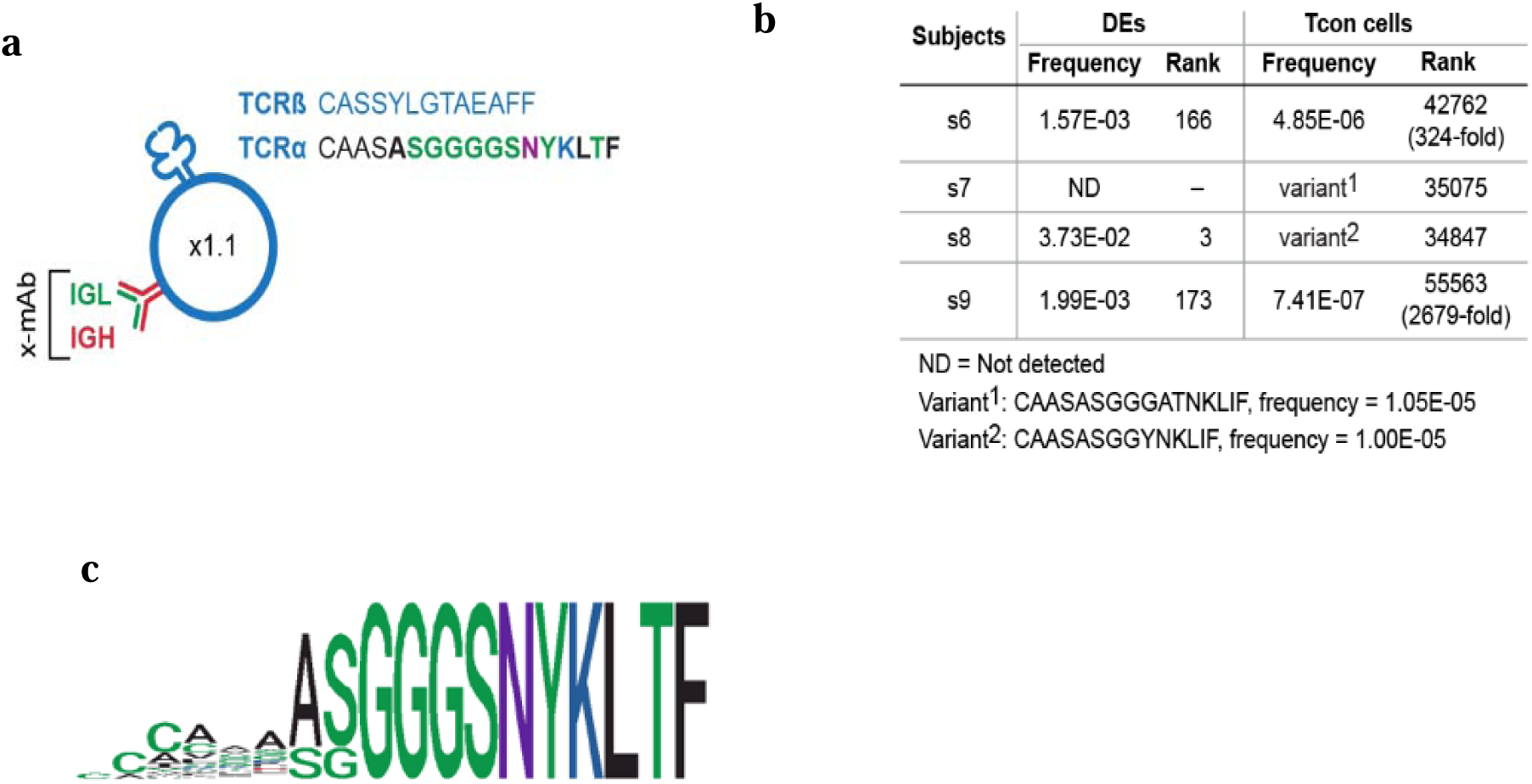
Association of public TCRα clonotypes by members of the TCRβ network. (**a**) Schematic shows CDR3 sequences of the TCRα and TCRβ-x chains of the TCR αβ coexpressed with the x-mAb in a DE lymphocyte clone. (**b**) Table shows frequency and rank of the TCRα-x chains in TCRα repertoires of DEs in indicated subjects. Fold change in the frequency of TCRα-x in DEs relative to Tcon cells in each subject is shown in parenthesis. (**c**) Weblogo shows the recurring CDR3 motif of the TCRα-x chain based on the sequence alignment consensus of retrieved sequences.

### An analogous network of BCR clonotypes is anchored by the x-mAb

Given that the x-mAb is the opposing partner of the TCRβ-x clonotype, we wondered whether the x-mAb is a member of a TCRβ-x-like family with characteristic invariant VHD or DJH motifs. Mining of the iReceptor and nPOD repositories did indeed identify the heavy chain of the x-mAb as a member of a large IGH family defined by a canonical CDR3 sequence (CARx(1-4)DTAMVYYFDYW), where x represents variable residues (**Fig. 6a**). Each clonotype is made of an invariant DJH motif recombined to one of various VH genes. We identified 7,937 clonotypes bearing the canonical CDR3 sequence in the iReceptor repository [from 274 repertoires, 24 labs and 103 subjects, 26 studies (**Supplementary Table 16a**)]. An additional 19 clonotypes bearing the canonical CDR3 sequence were detected in the nPOD database, 12 of them were from PLNs (**Supplementary Table 16b**). Of note, the TCRdb repository contains no IGH sequences. Figure 6b shows the common VDJ structure of the CDRH3 of the x-mAb, the prototype of the family (**Fig. 6b**). The invariant DJH (DTAMV<u>Y</u>YFDYW) motif (x-motif) is encoded by the DH5-18 (DTAMV) and JH04*01:01 (YFDYW) genes joined together by a palindromic tyrosine (underlined). The codon (TAT) for tyrosine is invariably generated by a p-mononucleotide (T) stitched to an ‘AT’ dinucleotide at the 5‣ recombining end of the JH04*01:01 gene (**Fig. 6c**). Participating VH genes as well as the JH04*01:01 gene used their full coding sequences with heterogeneity among family members limited to non-templated residues (N1-region) at the VH-DH boundary denoted by x in the canonical sequence (CARx(1-4)DTAMVYYFDYW). Intriguingly, a highly related sequence of the IGH-x CDR3 sequence (RQENFDTAMVYYF) is used as a variant surface glycoprotein (VSG), one of an array of potent antigens used by trypanosome brucei^14^. Thus, the x-mAb and its most enriched TCRβ partner belong to unique families of public antigen receptors made of V, D and J genes that are linked to one another in invariant motifs that are used to generate highly related CDR3 sequences.

**Fig. 6.**
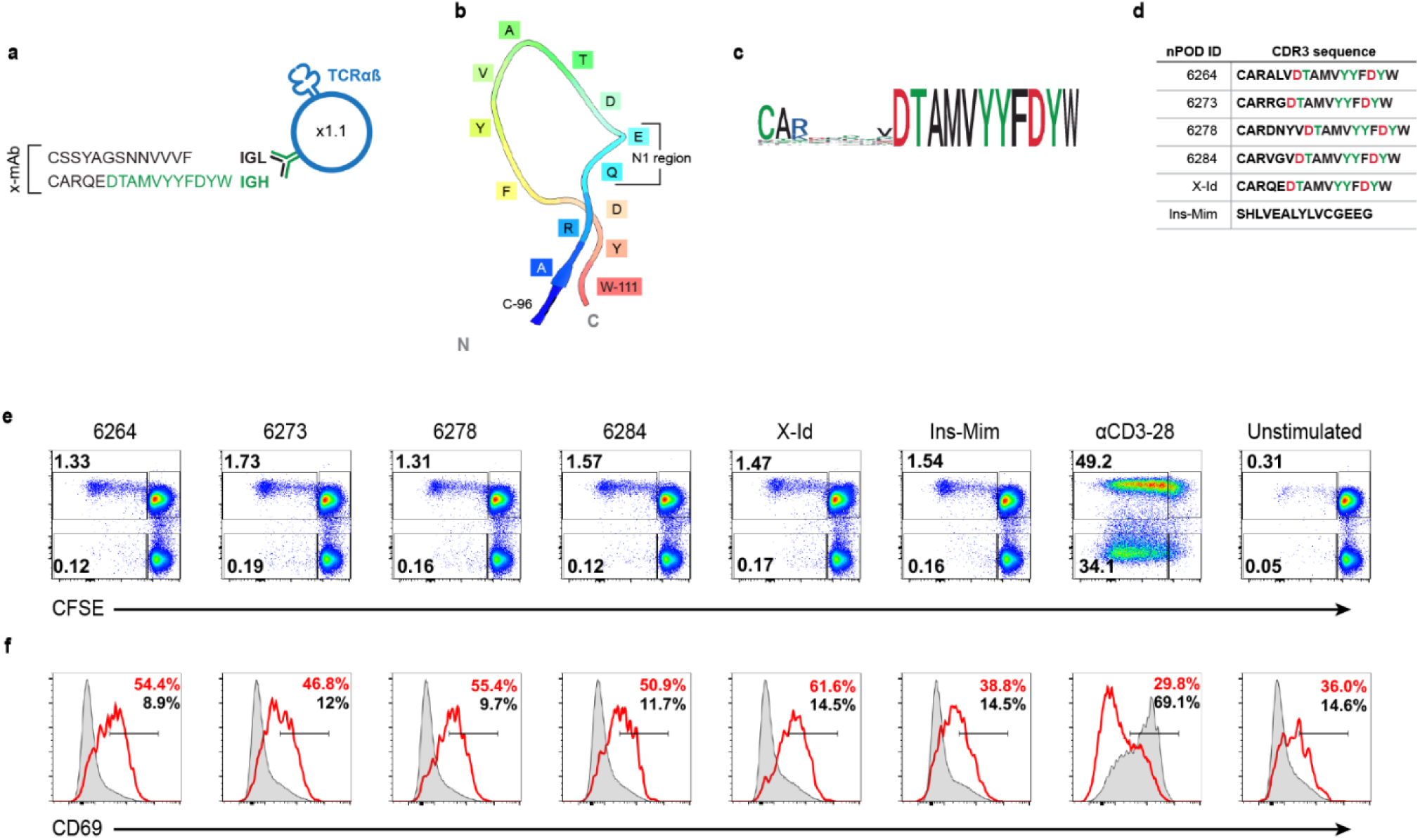
X-mAb is a prototype of the BCR network defined by signature CAR_x1-4_DTAMVYYFDYW sequence and immunogenic idiopeptides. The network members were generated by using a fixed DJH (DTAMVYYFDYW) motif with various VH genes. (**a**) Schematic shows that CDR3 sequences of IGL and IGH chains of the x-mAb coexpressed with TCRαβ in a DE clone. (**b**) Diagram shows the ribbon structure of the prototypic IGHV clonotype used by the x-mAb. The heterogeneity among network members is limited to the N1 region shown in bracket in the ribbon diagram of the IGHV clonotype of the x-mAb. (**c**) Weblogo shows the canonical CDR3 sequence of the IGH-x family based on the consensus of sequence alignment. Note heterogeneity limited to the N-region at the boundary with the shared invariant DJH motif. (**d**) Table shows CDR3 sequences of selected IGH-x clonotypes (source nPOD subjects) as well as the sequences of the x-Id of x-mAb and insulin mimotope that were used in the stimulation assay. (**e**) Dot plots show CFSE dilution by CD4 T cells from one T1D patients in response to stimulation with synthetic peptides of sequences depicted in the table D. Numbers indicate percentages. (**f**) Histogram overlays show CD69 upregulation by proliferated CFSE^low^ (red) and non-proliferated CFSE^hi^ (grey).

The x-mAb and an additional 50 clonotypes that were identical to the x-mAb except for lacking the (Q) residue at N1 region (CARQEDTAMVYYFDYW versus CAR*EDTAMVYYFDYW) were among the retrieved sequences (**Supplementary Table 16c**). These sequences were isolated from Bcon cells of healthy individuals, which is consistent with our published data showing that the x-mAb is rarely expressed among Bcon cells and that it is mainly expressed by DEs of T1D^4^. In addition, the lambda light chain that paired with the heavy chain of the x-mAb in DEs had 81 identical occurrences (CSSYAGSNNVVVF) in the iReceptor repository (**Supplementary Table 17**). Furthermore, more than 90% of the IGH-x family members, like the x-mAb, are of the IgM isotype, the most ancient antibody molecule in different vertebrates. The detection of x-mAb and large family of related clonotypes stands in stark contrast to the claim by Japp et. al. that they were not able to find the x-mAb or closely related sequence among Bcon cells or DEs of T1D subjects^15^. Thus, the heavy chain of the x-mAb of DEs is a prototype of a large family of IgM antibodies that is defined by a canonical CDR3 sequence and an invariant DJH motif.

### CDR3 sequences of the BCR network are immunogenic idiopeptides

Some immunoglobulin heavy chains V region peptides are immunogenic idiopeptides for T cells^16, 17^. In our hands, the x-idiotype (x-Id) of the x-mAb is an autoantigen and high-affinity ligand for the T1D-predisposing HLA-DQ8 molecule. The core sequence of the x-idiotype (<u>DTAMVYYFDYW</u>) is the denominator that defines the members of the BCR network. It is responsible for the high affinity binding to T1D-predisposing HLA-DQ8 molecule with the negatively charged aspartic residues occupying pocket 1 and pocket 9 of the DQ8 molecule and a pair of tyrosine in the core peptide^4^. On the other hand, no appreciable binding role was attributed to the heterogeneous residues at the N1 region, denoted by x in the signature sequence (CARx(1-4)DTAMVYYFDYW). In concordance, we predicted that the CDR3 idiotypes of clonotypes in networks encode antigenic T cell epitopes. We selected four clonotypes isolated from nPOD T1D subjects as representatives to test this possibility (**Fig. 6d**). MHC HLA binding analysis using the immune epitope database (IEDB) tool predicted strong binding of examined sequences to the DQ8 molecule. Subsequently, we directly examined the stimulatory ability of the peptides ex vivo using CD69 upregulation and CD4 T cell proliferation as readouts. We used PBMCs from a DQ8^+^ T1D subject as responders. All tested peptides stimulated CD4 T cells albeit with a varying degree of intensity (**Fig. 6e-f**). The results show that members of the BCR network not only share a signature CDR3 sequence, but also an antigenic functional property.

### The prototypes of TCR**αβ** and BCR networks are interacting partners

We used homology modelling and protein-protein docking to gain insights into the molecular interactions between the TCRβ-x clonotype and the x-mAb. The TCRαβ analyzed was made of the TCRα-x (CDR3: CAASASGGGGSNYKLTF) and TCR/β-x (CDR3: CASSYPGTAEAFF). The x-mAb was comprised of the heavy chain (CDR3: CARQEDTAMVYYFDYW) and the light chain (CDR3: CSSYAGSNNVVVF) that were cloned from BCRs of single DEs ^4^. We identified seven different x-mAb/TCRαβ complexes, and selected the three most stable modes, denoted as Complex 1, 3, and 6, respectively (**Fig. 7** and **Extended Data 8-9**) for further analysis. The selection was based on structural stability, contact areas and interaction energies between the x-mAb and the TCRαβ from at least 200 ns unbiased molecular dynamic (MD) simulations (**Extended Data 10**). The three complexes were subjected to umbrella sampling simulations to quantify the binding free energies (BFE) between the molecules (**Fig. 7a**). Among the three binding modes, Complex 6 was found to be the most stable with a BFE of -9.5 kcal/mol compared to -6.15 kcal/mol and -3.51 kcal/mol for Complex 3 and Complex 1, respectively (**Fig. 7b** and **Extended Data 9a**). In Complex 6, two CDR loop regions from the TCRα chain were involved in the binding: Met29 from the CDR1α (NS<u>M</u>FDY) and residues Ser95 to Ser100 (underlined) from the CDR3α (CAASA<u>SGGGGS</u>NYKLTF) (**Fig. 7c**). Two loop regions from the TCRβ chain also participated in the binding to the x-mAb. Residues Tyr95 to Gly97 (underlined) from the CDR3β (CASS<u>YPG</u>TAEAFF) were observed in both Complex 1 and Complex 3, whereas residues N28 to E30 from CDR1β (M<u>N</u>HEY) were only observed in Complex 1 (**Fig. 7d** and **Extended Data 9b-e**). Interestingly, the binding interface of all three complexes involved the CDR3 loop region SGGGGS, making it the common “paratope” for the most ranked x-mAb/TCR complexes as determined by our molecular modeling and simulations. Moreover, only Complex 6 exhibited a canonical tip-to-tip binding of the x-mAb and TCRαβ, consistent with being the complex with highest binding affinity. The interactions between the x-mAb as the prototype of the BCR network with a prototypic variant of the TCRβ network underscore a functional relationship between the two networks.

**Fig. 7.**
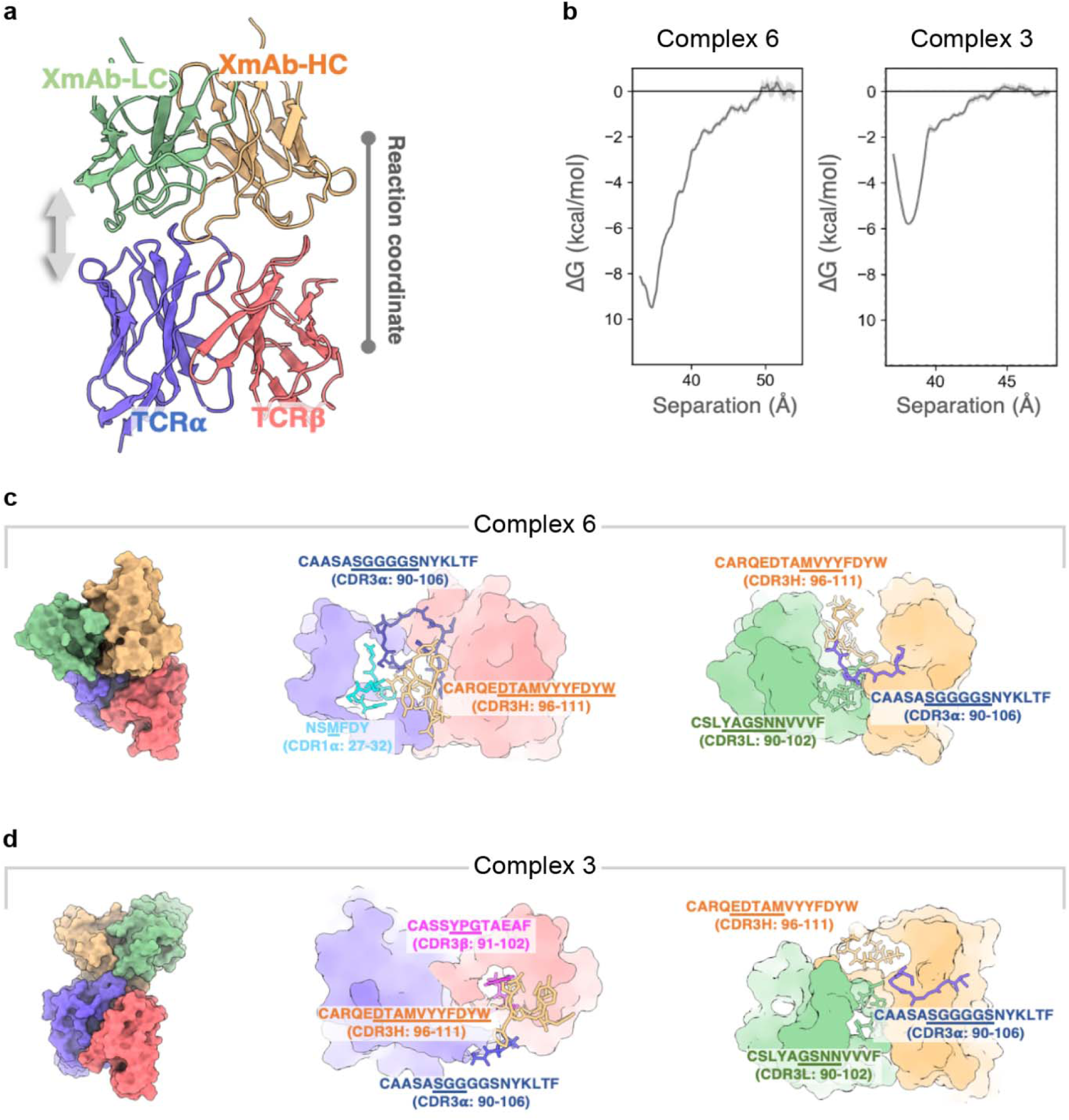
The prototypes of TCRαβ and BCR networks are interacting partners. (**a**) Schematics of the free energy calculation setup of the TCRαβ/x-mAb complex. The light and heavy chains of the x-mAb are colored in tan and green, respectively. The alpha (α) and beta (β) chains of the TCR are colored in blue and red, respectively. (**b**) The binding free energies for Complex 6 and Complex 3. (**c-d**) The overall structures and detailed molecular interactions for Complex 6 and Complex 3. **Left**, The overall structure of the TCR/x-mAb complex. **Middle**, CDR loops from the TCR, which interact directly with the CDR3H of the x-mAb, are specifically colored and labelled. Residues which are directly in contact with CDR3H are underlined. **Right**, CDR loops from the x-mAb, which interact directly with the CDR3α of the TCR, are specifically colored and labelled. Residues which are directly in contact with CDR3α are underlined.

## DISCUSSION

In this study, we identified and defined two networks of millions of TCRs and BCRs that are made from the CDR3 sequences of one TCRβ clonotype (CASSPGTEAFF) and one IGH (CARQEDTAMVYYFDYW) clonotype, respectively. The TCRβ network is made of clonotypes bearing the prototype CDR3 sequence (CASSPGTEAFF) or one of two signature CASSPGT-Jβx and Vβx-PGTEAFF (x means any J or V gene) sequences. The latter two are generated by splitting the VDJ joining of the prototypic CDR3 sequence into two overlapping VβD-(CASSPGT-) and -DJβ (-PGTEAFF) motifs that are used in highly recurring manners with various Jβs and Vβ genes, respectively. The IGH network is made of few clonotypes bearing the prototypic sequence (CARQEDTAMVYYFDYW) and thousands defined by a signature sequence (CARx_1-4_<u>DTAMVYYFDYW</u>) made from the DJH motif (underline) of the heavy chain of the prototypic IGH clonotype in combination with various VH genes. In total, we identified more than 8 million clonotypes that belong to the TCRβs and IGH networks. Remarkably, the prototypic TCRβ clonotype alone is associated with more than 63 diseases and conditions (cancers, infections, vaccinations, autoimmune diseases) indicating involvement in various immune and autoimmune responses.

In disregard to MHC restriction and consistent with the global usage, the prototypic TCRβ clonotype is used by CD4 and CD8 T cells encompassing memory, effector and regulatory subpopulations in numerous independent studies. Furthermore, the TCRβ-x clonotype is retrieved not from one or two panels that but from seven panels that recognize different COVID-19 epitopes with high confidence, indicating antigen recognition degeneracy. In addition, the CASSPGT sequence and its variants are used by many IGH clonotypes and by TCRβs of phylogenetically divergent species (rats, mice and pigs). Moreover, both the CASSPGT and PGTEAFF motifs are found in linear sequences of germline-encoded proteins (mainly enzymes) in humans and prokaryotes, indicating they are evolutionarily conserved. Repeated and versatile usage of the same motifs of the two networks indicate that it is unlikely random and must be for perhaps ensuring generation of antigen receptors bearing the ability to recognize important antigenic specificities.

Indeed, the members of the two families are apparently teleogically related. The strongest indirect evidence is coexpression of a member of the TCRβ network and prototypic IGH on dual expresser (DE) lymphocytes. Most direct evidence is provided by molecular simulations showing that x-mAb and TCR expressed on DEs interacts with each other at multiple paratopes. The interactions occur mainly through the invariant motifs. At the primary sequence level, the DJH motif encodes self-reactivity as indicated by the ability of synthetic peptides to stimulate a subset of CD4 T cells. Thus, the adaptive immune system appears highly committed to preserving these two prototypic sequences and their variants perhaps for a fundamental role that is too critical to leave for random VDJ recombination. Analysis and tracking usage of clonotypes of the two networks in various response and normal repertoires could lead to new hypotheses and mechanistic studies to accelerate understanding autoimmune and immune response repertoires of T and B cells.

Future studies should examine the mechanisms underlying the generation of the members of the networks. While there is no precedence to our knowledge of the systemic and variegated usage of the VDJ joining of the TCRβ, there is ample evidence of repeated usage of DJH-motifs with different VHs. One mechanism is VH replacement, a process whereby a VH gene of a fully assembled heavy chain is replaced by an upstream non-rearranged VH gene producing a new clonotype that retains the original DJH motif^18^. VH replacement, however, is a mechanism for antibody diversification that operates at the individual clonotype level to replace unproductive clonotypes by new productive clonotypes^19, 20^. An alternative, but not mutually exclusive, possibility draws on the fact that nearly half of the heavy chain genes in elasmobranch are completely or partially prejoined in the germ-line DNA, producing highly similar clonotypes using the fused VDH or DH motifs among different individuals^21, 22^. Of interest in this regard is the presence of a sequence highly similar to the canonical CDR3 that includes the x-motif sequence in the variant surface glycoproteins (VSG) of the African trypanosome parasite protozoan^23^. The presence of the TCRβ-x invariant motifs in germline genes of prokaryotes reinforces this possibility, but we cannot formally exclude random gene rearrangements as the mechanism behind the generation of the IGH-x family members. The results provide invaluable insights that could lead to new hypotheses and methods to track adaptive and pathogenic immune responses.

## Supporting information

All Extended Data 1-10

## MATERIALS AND METHODS

### Study design

The aim of this study was to characterize T cells recognized by the germline-encoded x-mAb in patients with T1D. We determined whether x-mAb-reactive T cells are phenotypically and transcriptionally distinct from non-reactive T cells (Tcon), their TCRβ repertoire and reactivity against insulin, the primary autoantigen in T1D. We also used molecular simulations to analyze interactions of the x-mAb with the public TCRαβ that was most enriched among reactive T cells. We assessed the frequency x-mAb-reactive T cells in randomly assigned T1D patients and in Healthy subjects (HC). Beyond analysis of cell frequency is we limited our study to PBMCs from T1D due to paucity of x-mAb-reactive T cells in HCs. Multicolor flow cytometry staining panels were used to compare surface phenotypes, cytokines of x-mAb-reactive T cells as compared to autologous Tcon cells in T1D patients. Further, x-mAb-reactive T cells and autologous Tcon cells were sorted in two batches and their transcriptional profile determined in parallel using bulk RNA-seq. We also sorted and compared TCRβ clonotype repertoires of x-mAb-reactive T and autologous Tcon cells using the ImmunoSEQ assay at the survey level. For *ex-vivo* assessment of immunogenic abilities of synthetic peptides representing the CDR3 sequences of selected clonotypes and PBMCs from a DQ8^+^ T1D donor were used as responders and CD69 upregulation and CFSE assay to measure to measure responses. T cell activation was performed after seven day of stimulation. Unstimulated cultures were used negative control and anti-CD3/CD28 stimulation as positive control. All data points and n values reflect biological replicates, and numbers of human donors are included, with the statistical test performed, in the caption for each figure.

### Human participants and PBMC isolation

All research was performed according to a protocol approved by the Johns Hopkins Institutional Board Review and conducted in accordance with the principles of Declaration of Helsinki. Written informed consent/assent was obtained from each participant. Patients with type 1 diabetes (T1D) that qualify under the American Diabetes Association criteria for classification were recruited at the Johns Hopkins Diabetes Center and Johns Hopkins Children’s Center. Donors with no previous T1D history were classified as healthy control subjects (HCs). Autoantibody profiling and HLA haplotypes (HLA genotype) of selected numbers of donors were performed at the Barbara Davis Center Autoantibody / HLA Core Laboratory in Denver. Donor demographics, including islet autoantibodies and HLA genotypes (s1-s9) are shown (**Supplementary Table 1**). Peripheral blood mononuclear cells (PBMCs) were isolated from heparinized blood by density gradient centrifugation using Ficoll-Paque (GE Healthcare) according to the manufacturer’s protocol. Isolated PBMCs were used fresh or aliquoted and kept frozen at −80L°C until used.

### Production and purification of the x-mAb

A previously generated EBV-immortalized dual expresser (DE) lymphoblastoid cell line (clone x1.1) was used as the source of the x-mAb^4^. Purification of x-mAb was performed using human IgM purification kit according to the manufacturer’s instructions (LigaTrap®), verified using SDS-PAGE, and the protein concentration determined using EZQ Protein Quantitation Kit (Invitrogen).

### Flow Cytometry

Fresh or rapidly thawed cryopreserved PBMCs were resuspended in complete tissue culture medium, washed with PBS and stained using established methods^4, 24^. Cell surface staining was performed on ice using predetermined optimal concentrations of indicated fluorochrome-conjugated antibodies.

Single-color-stained and fluorescence minus ones (FMO) were used for setting flow panel compensations. Samples were acquired by flow cytometry using Attune Nxt Flow Cytometer (ThermoFisher). Acquired samples (0.5 - 1 × 10^6^ events) were analyzed by FlowJo (TreeStar). Cell doublets were excluded in a step-wise manner from analysis using FSC-H vs FSC-W plots and SSC-H vs SSC-W (**Extended Data 1 and 3**). Specificity controls (human FcR blocking reagent (Miltenyi Biotec), FMO, dump gating, and isotype controls) were used as needed. When applicable, irrelevant cell types were used as internal biological controls and in the case of in-vitro cell culture stimulation assays, unstimulated samples served as negative controls.

### Detection of x-mAb-reactive T cells

Purified x-mAb was used for staining T cells in two consecutive steps. First, PBMCs were incubated with x-mAb (5 μg) for 1 h at 4 ^°^C and washed twice with FACS buffer. Next, pelleted cells were resuspended in FACS buffer and incubated for 50 mins with AF-488 or FITC-conjugated anti-human IgM secondary antibody with a cocktail of fluorochrome-conjugated antibodies to detect indicated cell markers. Samples were washed twice with FACS buffer and acquired by Attune Nxt flow cytometry and analyzed using the FlowJo software. In limited experiments, directly conjugated x-mAb (AF-488-x-mAb) was used. Briefly, Alexa Fluor amine-reactive dye AF-488-TFP (Molecular probes / Invitrogen) was prepared by dissolving 1 mg in 100 μL of DMSO and used to label x-mAb according to manufacturer’s instruction. X-mAb in PBS was added to the AF-488-TFP, mixed well and incubated at room temperature for 1 h. Unconjugated dye was removed by dialysis at 2–6 °C overnight with four buffer changes. AF-488-x-mAb was used for staining PBMCs together with the cocktails of indicated antibodies. To exclude x-mAb binding to Fcα/uR, samples were preincubated with human serum for 30 mins before staining with AF-488-x-mAb. T cells that stained with x-mAb are referred to Tx cells and those negative for x-mAb staining as Tcon cells.

### Comparative phenotypic and functional analysis of Tx and Tcon cells

Samples were stained with relevant antibodies and x-mAb-reactive and non-reactive Tcon cells gated and analyzed for: (1) Indicated activation markers. (2) Different T cell subsets: Naïve T cells, T_N_ (CD45RA^+^CCR7^+^), effector memory cells re-expressing CD45RA, T_TEMRA_ (CD45RA^+^CCR7^-^), effector memory, T_EM_ (CD45RA^−^CCR7^−^), and central memory, T_CM_ (CD45RA^−^CCR7^+^). Treg cells were defined as TCRαβ^+^CD4^+^Foxp3^+^CD25^+^, whereas DEs are defined as TCRαβ^+^IgD^+^IgM^+^. For detection of Treg cells, samples were surface-stained, fixed and permeabilized using Foxp3 transcription factor staining buffer kit (eBioscience) before intracellular with anti-Foxp3 antibody at room temperature. (3) Cytokines, which were detected using intracellular staining after stimulation of samples with phorbol 12-myristate 13-acetate (PMA) (50 ng/mL) and ionomycin (500 ng/mL) in the presence of Golgi-Plug (2 mmol/L) in CTM for 4-5 h at 37 ^°^C in a humidified 5% CO_2_ incubator using established protocols^4^.

### Sorting of peripheral T cells into x-mAb-reactive T cells and non-reactive Tcon cells

Sorting of PBMCs into x-mAb-reactive and Tcon cells was used for TCR repertoire (TCRβ and TCRα) and transcriptome analysis. PBMCs were stained with the x-mAb followed by staining with anti-TCRαβ and CD4 antibodies as described above and sorted using MoFlow Legacy high-speed cell sorter (**Extended Data 5**). Propidium iodide (PI) was added immediately prior to sample acquisition to identify and exclude dead cells. x-mAb-reactive and Tcon cells were gated and sorted cells in RPMI media containing 50% FBS . Numbers of acquired x-mAb-reactive CD4 T cells range from 8,000 to 30,000 cells and an equivalent number of Tcon cells were collected per respective sort.

### TCR repertoire analysis

Sorted Tx and Tcon cells were washed once with PBS and total genomic DNAs were directly extracted using a QIAmp DNA blood mini kit according to the manufacturer’s instructions (Qiagen). Extracted DNA samples were quantified and shipped to Adaptive Biotechnologies for TCRBV and TCRAV sequencing using the ImmunoSeq platform at the survey level resolution. The ImmunoSeq assay combines multiplex PCR with high-throughput sequencing and sophisticated bioinformatics pipeline for CDR3 region analysis10-12. Raw ImmunoSeq data were processed with ImmunoSeq Analyzer 2.0 software (Adaptive Biotech) and saved at the website of Adaptive Biotechnologies. Various VDJ combinations used by different clonotypes can be visualized by using CDR3 sequences and the IMGT/V-QUEST algorithm.

For our own analysis, measurement metrics of processed data were exported in .tsv file format and analyzed using the R platform. Clones of uncertain vGene identity or out-of-frame were excluded from downstream analysis. For the vGene for each cell type in the individual samples, the counts of distinct cell clones were obtained by summing the metric of the “estimated number of cell genomes present in the sample”, upon which the corresponding percentages were calculated. The percentage quantification provides a uniform basis for the vGene (VH and Vβ) usages to be fairly and consistently compared across the different cell types and samples, minimizing any effects that could result from sequencing differences and low x-mAb-reactive T cell numbers. Percentages were visualized with bar plots to make straight comparisons of vGene usages between different cell types. The presence or absence of vGenes was determined on the basis of the vGene usages.

### Assessing stimulatory capacity of x-IGH family CDR3 peptides

We selected CDR3 sequences from the IGH-x clonotypes retrieved from the nPOD database (**see** Fig. 6d) as representatives to assess their stimulatory ability *ex vivo* using CFSE proliferation assay. The peptides were custom-synthesized with >95% purity (Peptide 2.0) and stock solution (2.0 mg/ml) were prepared by dissolving freeze-dried peptides in sterile water and kept at -80 ^°^C until use. CFSE-labeled PBMCs from a DQ8^+^ subject were cultured into wells of 24-well tissue culture plate (1.5 - 2.0 × 10^6^ cells / well in 1 mL CTM) and stimulated in the presence of absence of different peptides (10 μM) at 37 ^°^C, 5% CO_2_ incubator for 7 days. Cultures stimulated with anti-CD3-CD28 antibodies were used as positive controls. Cultures were harvested and surface stained for indicated markers, acquired by FACS and analyzed for CFSE dilution using FlowJo software. The frequency of CFSE^low^ proliferated T cells was determined for each peptide. We also used upregulation of the CD69 activation as a second readout of T cell activation.

### Homology modelling and protein-protein docking

The amino acid sequences of TCRαβ of x-mAb were obtained from our previous work^4^. The 3D structures of TCRαβ and x-mAb were generated using Modeller ver. 9.22^25^. One-hundred models of the TCRαβ/x-mAb complexes with different binding orientations were generated using GRAMM-X webserver^26^. Seven models were chosen as initial structures for MD simulation, in which at least one residue in the CDR3 of the heavy chain of x-mAb (DTAMVYYFD, residues 101-109) was found to be within 6 Å from the CDR1-3 (residues 26-31, 50-55, and 93-103) of both α and β chains of the TCR (**Extended Data 10**).

### Molecular dynamics (MD) simulations

We solvated the systems with explicit TIP3P water molecules [10.1063/1.445869]. In order to mimic the physiological environment, we added appropriate sodium and chloride ions to neutralize the system at 150 mM ionic concentration. Then, the energy minimization, NVT equilibrium, NPT equilibrium, and 200 ns MD production run, with 2 fs timesteps, were sequentially performed. The temperature was maintained at 300 K by v-rescale method [10.1063/1.2408420], and the pressure was coupled by Parrinello-Rahman method [10.1063/1.328693] to keep the pressure at 1 atm. For long-range electrostatic interaction, we used particle mesh Ewald (PME) method [10.1063/1.464397] with cut-off distance set to 12 Å. Meanwhile, the cut-off distance of Van der Waals (VdW) interaction was also set to 12 Å. In addition, we used Verlet scheme for neighbor searching, LINCS algorithm to deal with bond constraints and converted all bonds containing hydrogen atoms into constraints [10.1002/(SICI)1096-987X(199709)18:12<1463: AID-JCC4>3.0.CO;2-H]. All simulations were performed using GROMACS [10.1016/j.softx.2015.06.001] and CHARMM36 force field [10.1021/ct300400x] [10.1038/nmeth.4067].

### Umbrella sampling

The binding free energy was calculated using the steered MD with umbrella sampling. We first performed the steered MD simulations to pull apart the x-mAb/TCR complexes and used the distance between the center of mass of the TCR and the center of mass of the x-mAb as the reaction coordinates (Fig. 7a). We defined the spacing of the windows with every 0.02 nm to 0.05 nm for umbrella sampling to obtain the binding free energy. We calculated the potential of mean force using the WHAM software [Grossfield, Alan, “WHAM: the weighted histogram analysis method”, version 2.0.10, http://membrane.urmc.rochester.edu/wordpress/?page_id=126] and estimated the errors using bootstrapping analysis. All simulations performed at 300 K.

### Statistical analysis

Statistical significance of the results was performed using Prism 9 (GraphPad Software). Analysis was performed using independent samples t-test or a paired sample student t-test as appropriate. p≤0.05 was considered as statistically significant. The results were expressed as the mean ± SEM or median and range, as appropriate.

**Acknowledgments:** We thank participants enrolled in this study and their families for their blood donation. This work was supported by the W.M. Keck foundation, NIH, United States grant R0 AI099027 (A.R.A.H.); the Norman Raab Foundation; and Diabetes Research Connection grant (R.A.).

**Author contributions:** R.A. designed and performed experiments, analyzed data, prepared figures and tables. N.M. performed experiments, analyzed data, and prepared figures. N.M. retrieved sequences from public databases, organized sequences and prepared supplementary tables and figures. R.A. and N.M. contributed equally to the manuscript. R.A.H. and P.P. designed and performed experiments. H.Z. and J.M performed sorting of Tx and conventional T cells. K.C.C., Y. S, D.R.B, S.L, and R.Z. designed, performed, analyzed, wrote, and interpreted the MDS simulations. A.G., J.H., and R.M.W. recruited donors. C.J. did differential gene expression and reconstruction, prepared heatmaps. C.J. performed statistical analysis of immuno-seq data. T.D. recruited donors and discussed, critically reviewed, and edited the manuscript. A.R.A.H. oversaw the study, identified the TCRβ-x and IGH-x families and wrote the original manuscript. All authors critically reviewed and edited the manuscript.

**Competing interest:** The authors declare no competing interests.

**Data and materials availability:** All data, code, and materials used in the analysis will be available from the corresponding authors upon reasonable request according to NIH and Hopkins data sharing policy. The raw RNA-seq data generated from this study will be deposited at NCBI database and available from Gene. TCR repertoires will be available at Adaptive Biotech Analyzer.

## Supplementary Figures

-Extended Data 1 – Gating strategy to identify x-mAb-reactive T cells using Attune Nxt flow cytometer
-Extended Data 2 – Identification of x-mAb-reactive CD8 T cells
-Extended Data 3 – X-mAb interaction with T cells is not mediated by FC mu receptor for IgM (FCμR)
-Extended Data 4 – Production of cytokines by x-mAb-reactive CD4 and CD8 T cells
-Extended Data 5 – Gating strategy used for sorting of x-mAb reactive and non-reactive autologous CD4 Tcon cells using MoFlow sorter
-Extended Data 6 – The TCRβs clonotype is widely used in various diseases and exhibited degenerate antigen recognition
-Extended Data 7 – Tissue distribution and usage of TCRβ-x clonotype (CASSPGTEAFF) by CD4 and CD8 T cells in nPOD subjects
-Extended Data 8 – Molecular surfaces of TCRαβ and x-mAb
-Extended Data 9 – Binding of the x-mA to TCRαβ in Complex 1
-Extended Data 10 – The flowchart of TCRαβ/x-mAb complex modelling

## Supplementary Tables

-Supplementary Table 1 – Patient/donor demographics
-Supplementary Table 2 – Tx and Tcon cells TCRβ repertoires for patients s1 through s5
-Supplementary Table 3 – TCRβ iReceptor data for CASSPGTEAFF
-Supplementary Table 4 – TCRdb data for CASSPGTEAFF
-Supplementary Table 5 – nPOD data for CASSPGTEAFF
-Supplementary Table 6a – TCRβ iReceptor data for CASSGPTEAFF
-Supplementary Table 6b – TCRdb data for CASSGPTEAFF
-Supplementary Table 6c – nPOD data for CASSGPTEAFF
-Supplementary Table 7 – nPOD data for CASSPGT
-Supplementary Table 8a – TCRβ iReceptor data for PGTEAFF
-Supplementary Table 8b – nPOD data for PGTEAFF
-Supplementary Table 9a – TCRβ iReceptor data for CASSGPT
-Supplementary Table 9b – nPOD data for CASSGPT
-Supplementary Table 10 – TCRβ iReceptor data for CASSPGI
-Supplementary Table 11 – TCRβ iReceptor data for CASSPGGGW
-Supplementary Table 12a – IGH iReceptor data for CASSPGT
-Supplementary Table 12b – IGH iReceptor data for CASSGPT
-Supplementary Table 12c – IGH iReceptor data for CASSPGI
-Supplementary Table 12d – IGH iReceptor data for CASSPGGGW
-Supplementary Table 13 – Additional examples of the CASSPGT and PGTEAFF in germline-encoded proteins
-Supplementary Table 14a – X and Tcon cells TCRα repertoires for patients s6 through s10
-Supplementary Table 14b – Tx and Tcon cells TCRα repertoires for patient s1
-Supplementary Table 15 – iReceptor data for ASGGGSNYKLTP, CAASGGGGNYKLTF, ASGGGGSNYKLTF, CAASASGGGNYKLTF, CAASASGGGGSNYKLTF sequences
-Supplementary Table 16a – IGH iReceptor data for DTAMVYYFDYW
-Supplementary Table 16b – nPOD data for DTAMVYYFDYW
-Supplementary Table 16c – iReceptor data for CAR*EDTAMVYYFDYW
-Supplementary Table 17 – iReceptor data for SSYAGSNNVVVF (lambda light chain of the x-mAb)

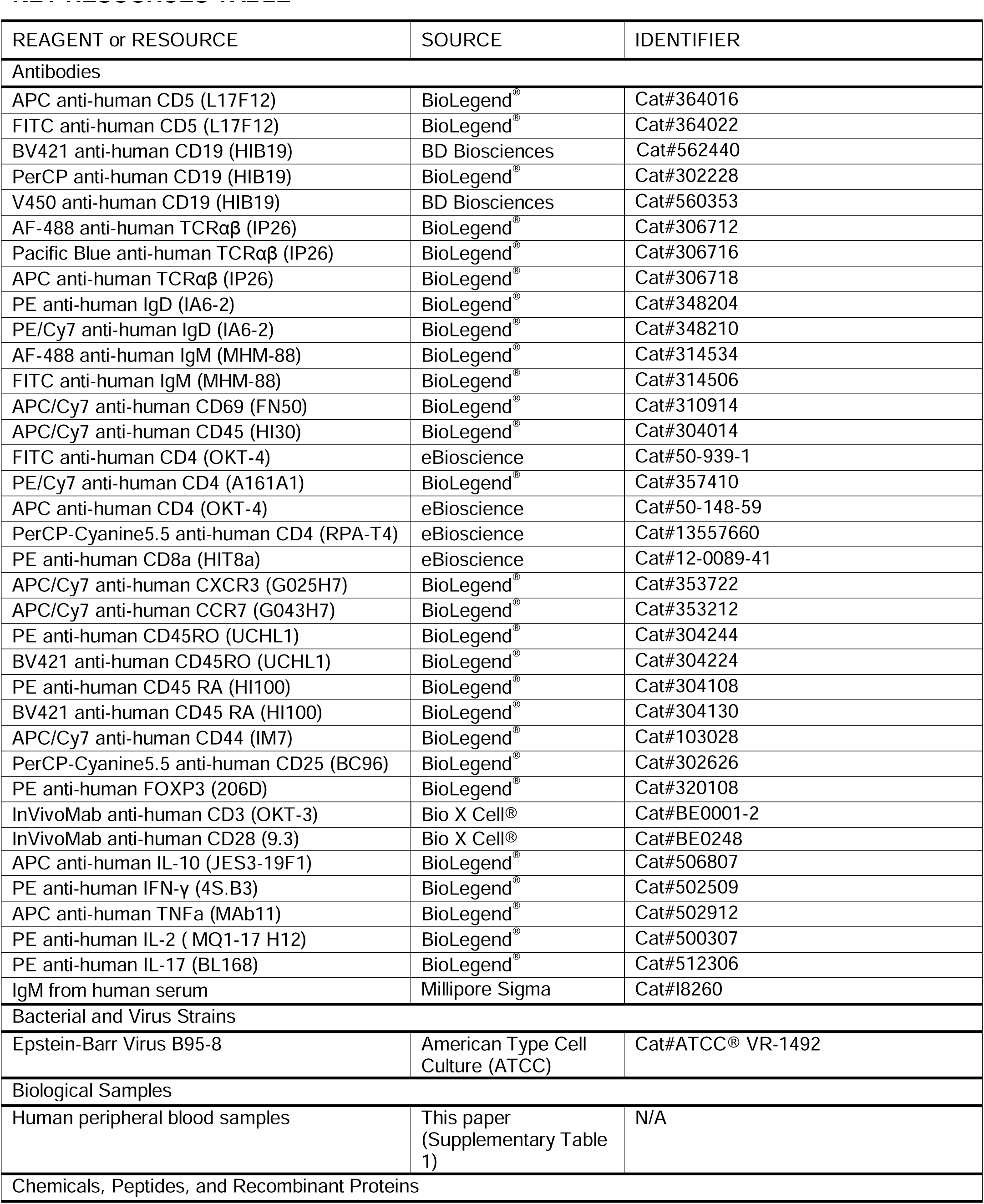
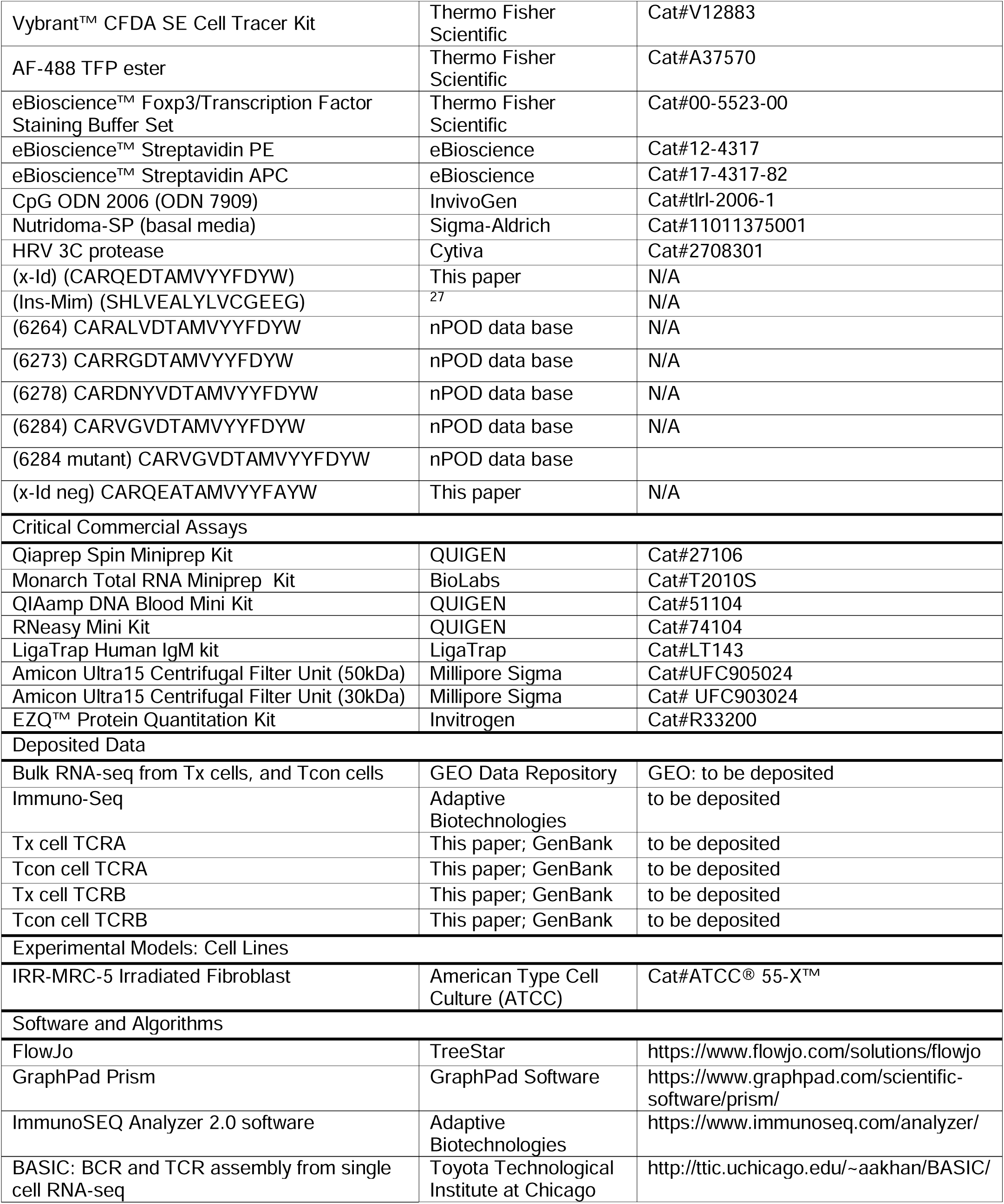
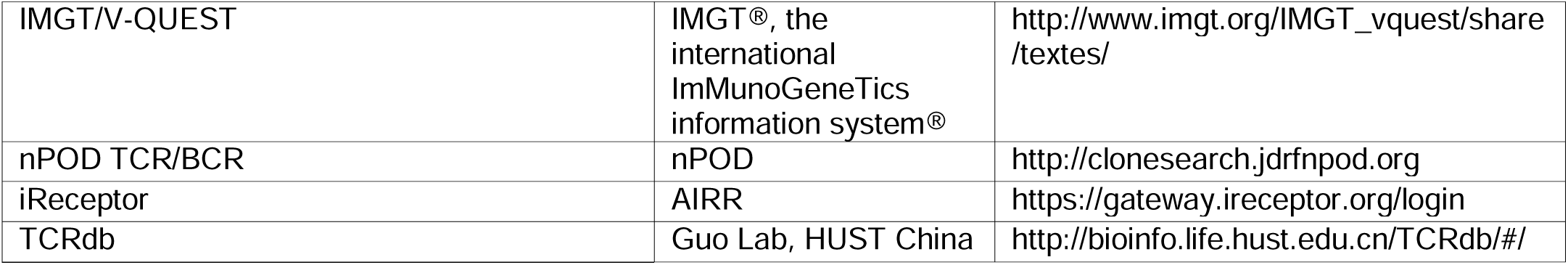
KEY RESOURCES TABLE

