## Supplementary material for "Apposed networks of interacting TCRs and BCRs exhibiting mosaicked CDR3 sequences made of fixed junctional motifs": All Extended Data 1-10

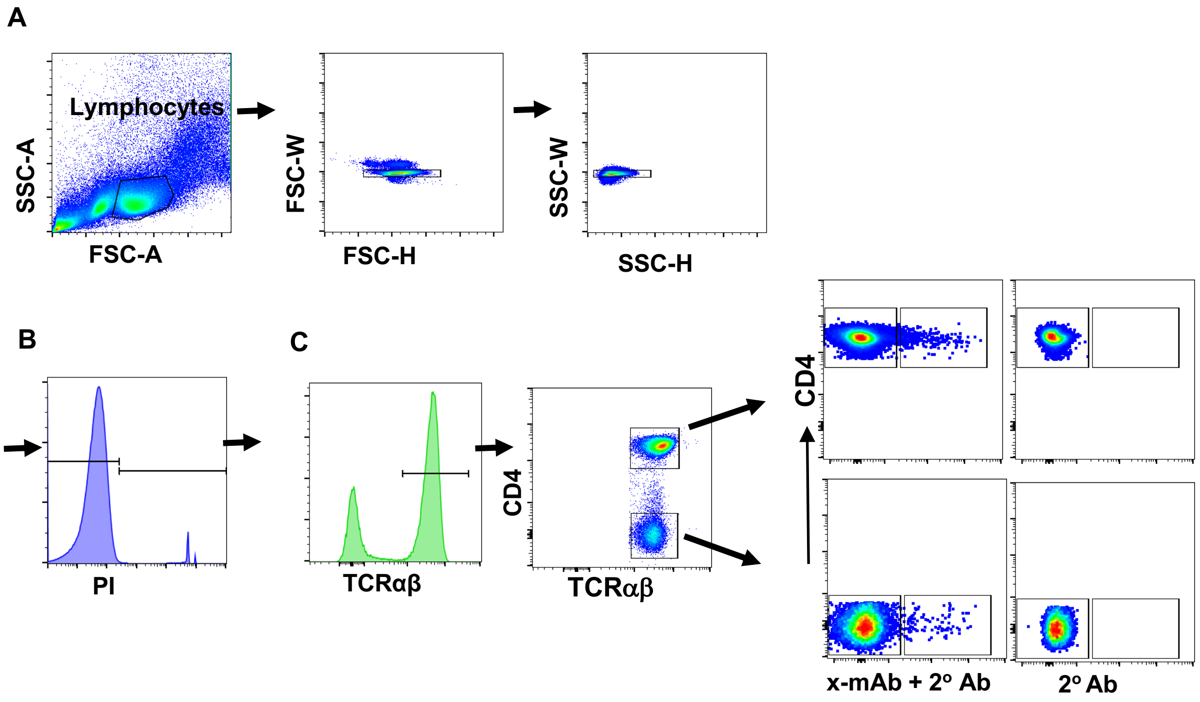


**Extended Data 1. Gating strategy to identify x-mAb-reactive T cells using Attune Nxt flow cytometer.** PBMCs were stained with unlabeled x-mAb followed by staining with fluorochrome-conjugated anti-TCRαβ, CD4 and anti-human IgM secondary antibody (2° Ab) as described in Materials and Methods. Single-color staining was used for compensation. Fluorescent-minus one (FMO) staining by anti-human IgM secondary antibody (2° Ab) in the absence of x-mAb was used to assess background staining. Acquired samples were analyzed by FlowJo software. (**A**) Lymphocytes were gated and doublets excluded using FSC-Height versus FSC-Width and SSC-Height versus SSC-Width plots. (**B**) Propidium Iodide (PI) was used to exclude dead cells. (**C**) TCRαβ$^+$ cells were gated and separated into CD4$^+$ and CD4$^-$ (i.e CD8$^+$) subsets and analyzed for x-mAb binding.

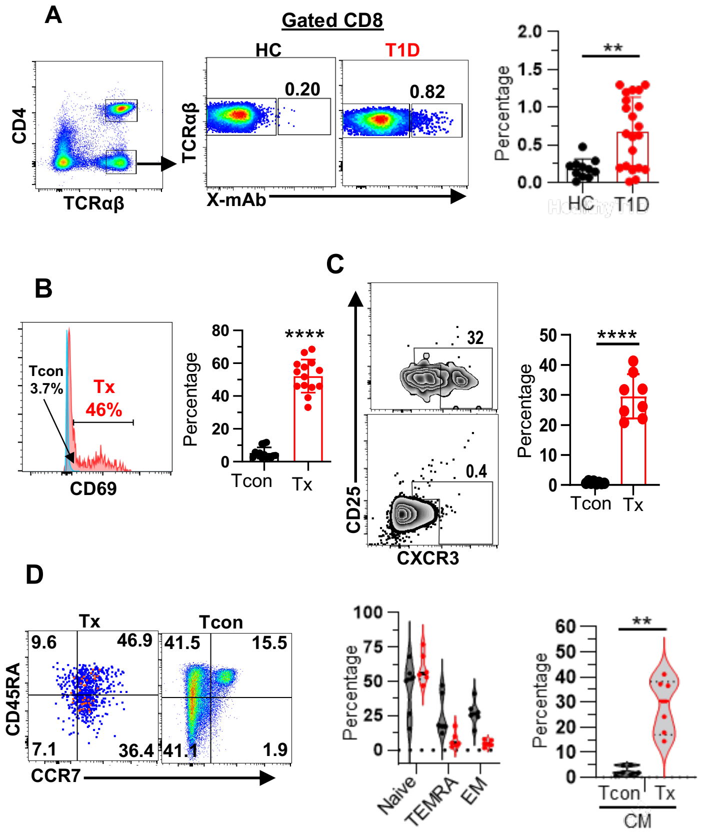


**Extended Data 2. Identification of x-mAb-reactive CD8 T cells. (A)** Representative dot plots depicting binding of a subset of CD8 T cells (identified as $CD4^-$ $TCR^+$) by x-mAb (Tx cells) in HC and T1D subjects. Numbers indicate percentages. **Graph** showing cumulative data (Mean ± SEM; T1D (n=21) and HC (n=10)). **P<0.01 by two samples independent t test. (**B**) **Histogram overlay** showing expression of CD69 by CD8 Tx (red) and Tcon (blue) cells. Numbers indicate percentages. **Graph** shows cumulative data (n=14 subjects). ****P<0.0001 by paired t test. **(C) Representative dot plots** showing expression CD25 and CXCR3 by CD8 Tx and Tcon cells. Numbers indicate percentages. **Graph** shows cumulative data (Mean ± SEM; n=8). ****P<0.0001 by paired t test. **(D)** Representative dot plots showing expression of CD45RA and CCR7 by CD8 Tx and Tcon cells. **Left**, violin graph shows cumulative data of naïve, TEMRA, and effector memory subsets among Tx and Tcon subsets. **Right**, violin graph shows cumulative data (Mean ± sem; n=6) for $T_{CM}$ central memory subset among CD8 Tx and Tcon subsets.

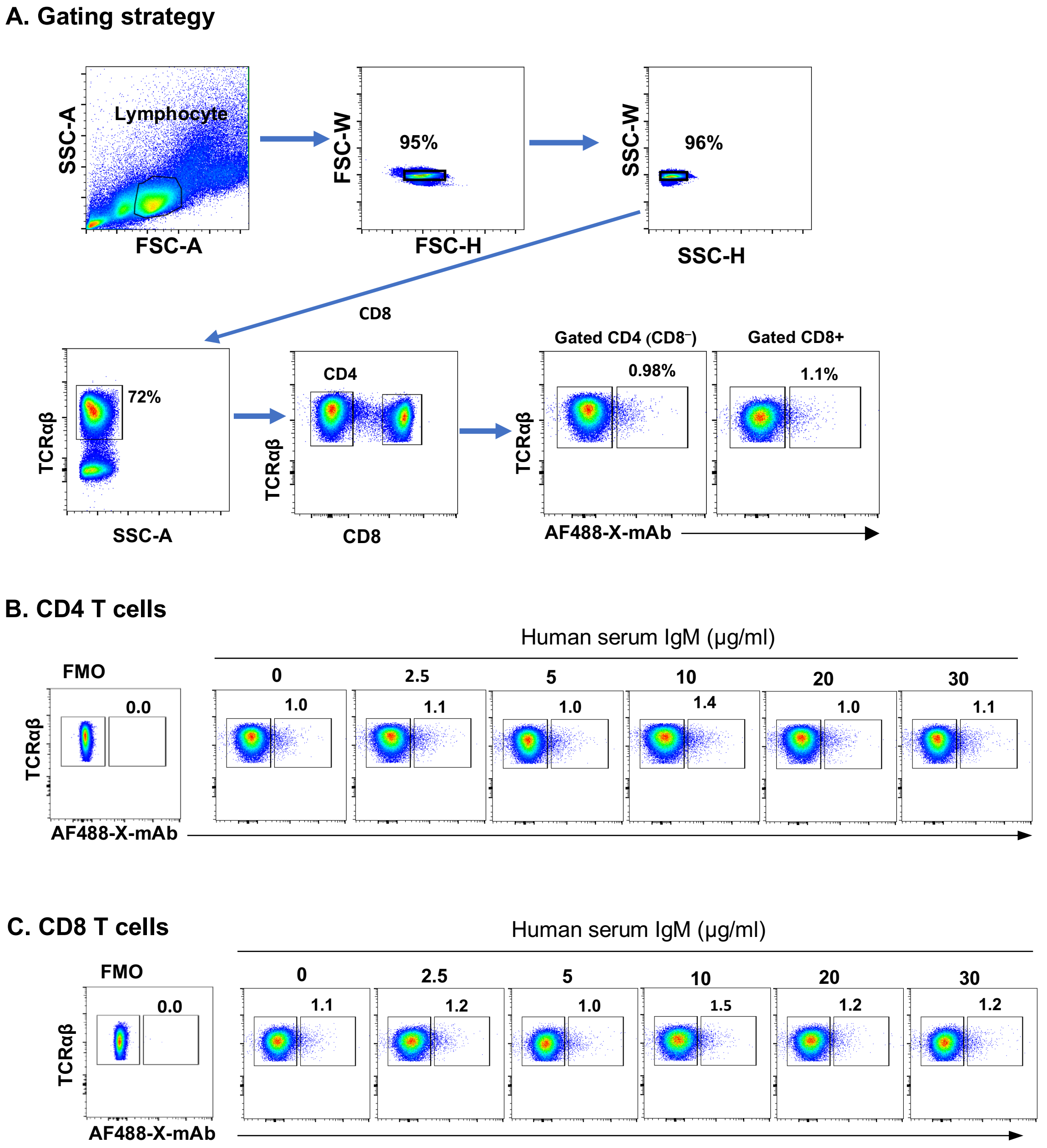


**Extended Data 3. X-mAb interaction with T cells is not mediated by FC mu receptor for IgM (FCµR).** To determine whether binding of the x-mAb to T cells is influenced by FCµR, PMBCs were preincubated with an increasing dose of human IgM serum for 30 minutes before staining with AF-448-labeled x-mAb followed by surface staining with anti-TCRαβ and CD8 antibodies as described in Materials and Methods. **(A) Dot plots** show sequential gating of stained PBMCs to exclude doublets and dead cells (used PI not shown), and to identify CD4 and CD8 T cell subsets. (**B-C**) **Left, FMO dot plots** show no background staining of CD4 (**A**) and CD8 (**B**) subsets of T cells in samples stained with anti-TCRαβ and CD4 antibodies in the absence of AF-488 x-mAb. **Right, dot plots** show percentages of CD4 (**B**) and CD8 T cells (**C**) that bound the x-mAb in samples pretreated with indicated concentration of human serum IgM. Note, CD4 T cells are identified as CD8- TCRαβ$^{+}$ cells.

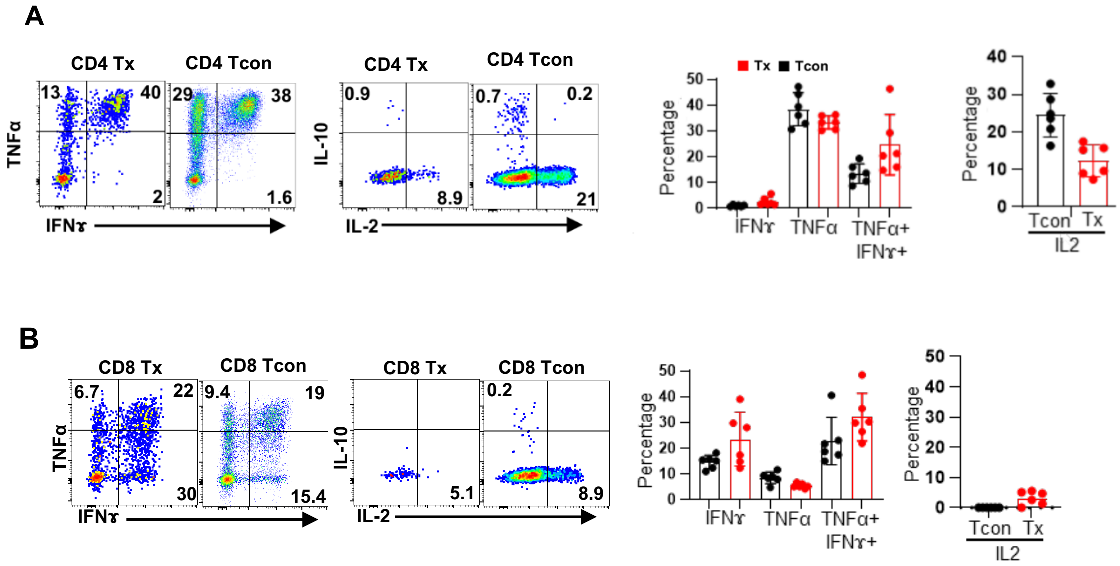


**Extended Data 4. Production of cytokines by x-mAb-reactive CD4 and CD8 T cells.** (**A**) Representative dot plots show intracellular expression of TNFα-vs-IFN-γ (left) and IL-10-vs-IL-2 (right) by gated CD4 Tx and Tcon cells. **Graphs** show cumulative data (n=6). IL-10 expression showed a negligible difference across the two subsets. (**B**) Representative dot plots show intracellular expression of TNFα-vs-IFN-γ (left) and IL-10-vs-IL-2 (right) by gated CD8 Tx and Tcon cells. **Graphs** show cumulative data (n=6). IL-10 expression showed a negligible difference across the two subsets.

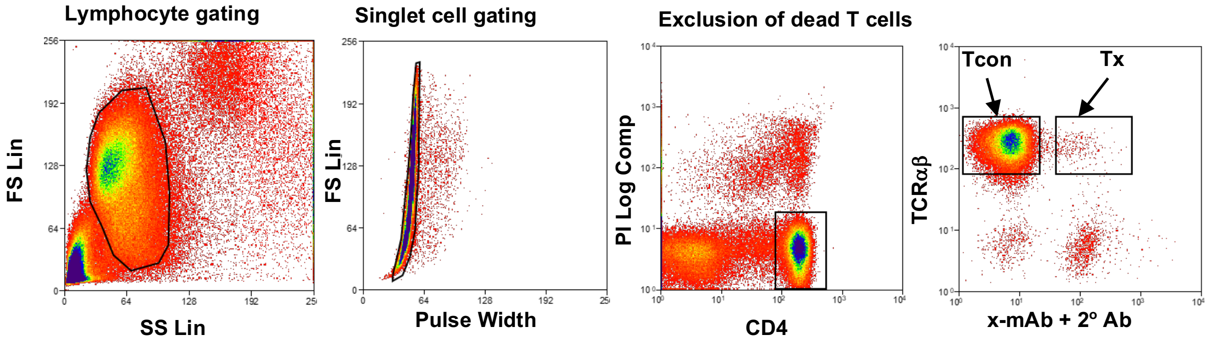


**Extended Data 5. Gating strategy used for sorting of x-mAb-reactive and non-reactive autologous CD4 Tcon cells using MoFlow sorter.** Dot plots show sequential gating of lymphocytes to exclude doublets and dead cells (using PI staining). Total live CD4 T cells were gated and sorted into x-mAb-reactive and non-reactive Tcon cells.

**A**

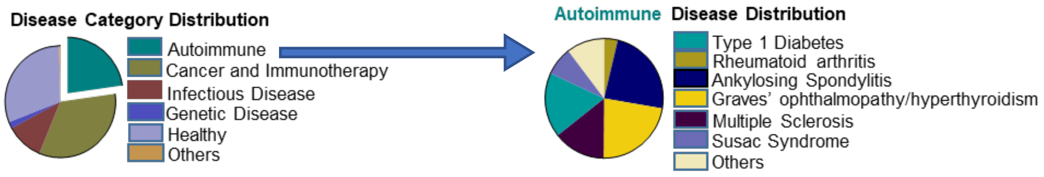


**B**

| TCR BioIdentity | Experiment | ORF Coverage | Amino Acids |
|---|---|---|---|
| CASSPGTEAFF+TCRBV04-02+TCRBJ01-01 | ePD87 | ORF1ab | ALRKVPTDNYITTY,KVPTDNYITTY |
| CASSPGTEAFF+TCRBV12-03+TCRBJ01-01 | eOX52 | ORF1ab | FVDGVPFVV |
| CASSPGTEAFF+TCRBV04-01+TCRBJ01-01 | eXL27 | ORF1ab | RQLLFVVEV |
| CASSPGTEAFF+TCRBV27-01+TCRBJ01-01 | eXL32 | envelope,ORF1ab | APAHISTI,LIVNSVLLFL,LLFLAFVVFL,SVLLFLAFV |
| CASSPGTEAFF+TCRBV09-01+TCRBJ01-01 | eOX52 | ORF1ab | YLNTLTLAV |
| CASSPGTEAFF+TCRBV07-09+TCRBJ01-01 | eQD112 | surface glycoprotein | DLPIGINITR,INITRFQTL,LPIGINITRF |
| CASSPGTEAFF+TCRBV07-09+TCRBJ01-01 | eMR17 | nucleocapsid phosphoprotein | SQASSRSSSR |

**Extended Data 6. The TCRβ-x clonotype is widely used in various diseases and exhibited degenerate antigen recognition**. (**A**) **Left Venn diagram** shows association of the TCRβ-x clonotype with different disease conditions as determined by mining the TCRdb repository. **Right Venn diagram** shows the top five autoimmune diseases associated with the TCRβ-x clonotype according to our search of the TCRdb database. (**B)** The TCRβ-x clonotype was identified among high-confidence SARS-CoV-2-specific TCRs specific for indicated individual or multiple peptides using Multiplex Identification of Antigen-Specific T-Cell Receptors Assay (MIRA). The information is derived from the ImmuneCODE MIRA Release 002 (ref. PMID: 32793896).

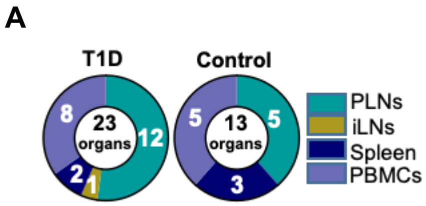


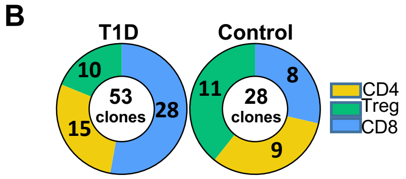


**C.** Tissue distribution and numbers of T cells bearing TCRβ-x clonotype in nPOD subjects

| Organ | **T1D** (n=23 subjects) | | | **Control** (n=7 subjects) | | |
|---|---|---|---|---|---|---|
| | Total CD4 | CD4 Treg | Total CD8 | Total CD4 | CD4 Treg | Total CD8 |
| **PLN** | 13 | 6 (46%) | 14 | 6 | 5 (83%) | 7 |
| **spleen** | 6 | 2 (33%) | 10 | 8 | 4 (50%) | 0 |
| **iLN** | 5 | 2 (40%) | 4 | 6 | 2 (33%) | 1 |

**Extended Data 7. Tissue distribution and usage of the TCRβ-x clonotype (CASSPGTEAFF) by CD4 and CD8 T cells in nPOD subjects**. (**A**) **Venn diagrams** show tissue distribution of the TCRβ-x clonotypes in 23 organs of 16 T1D subjects and 13 organs of 7 control nPOD subjects. (**B**) Venn diagrams show usage of the TCRβ-x clonotype by CD4 and CD8 T cells in T1D and control subjects. (**C**) **Table** shows total numbers and lineages of TCRβ-x-expressing T cells in different organs of the 23 nPOD T1D and 7 control subjects. Total CD4 T cells included Treg clones. Number of sequences isolated from Treg cells; their percentages out of total CD4 T cells are shown in parentheses. Only one clonotype was retrieved from PBMCs (not shown). Venn diagrams were generated using the TCR/BCR nPOD search program.

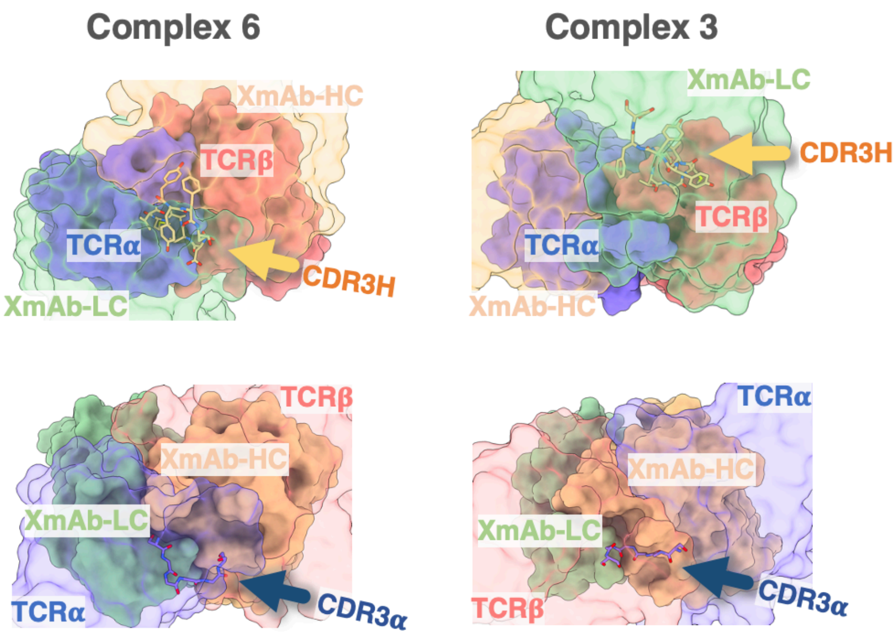


**Extended Data 8. Molecular surfaces of TCRαβ and x-mAb**. **Top panel**, top view of TCRαβ in Complex 6 and Complex 3. The TCRαβ is shown as solid surfaces with the α and β chains colored in blue and red, respectively. The x-mAb is shown as transparent surfaces with the light and heavy chains colored in green and tan, respectively. The CDR3H loop (DTAMVYYFD) of the x-mAb is explicitly shown as licorice. **Bottom panel**, top view of the x-mAb molecular surfaces in Complex 6 and Complex 3. The x-mAb is shown as solid surfaces with the light and heavy chains colored in green and tan, respectively. The TCRαβ is shown as transparent surfaces with the α and β chains colored in blue and red, respectively. The CDR3α loop (CAASASGGGGSNYKLTF) of the TCRαβ is explicitly shown as licorice.

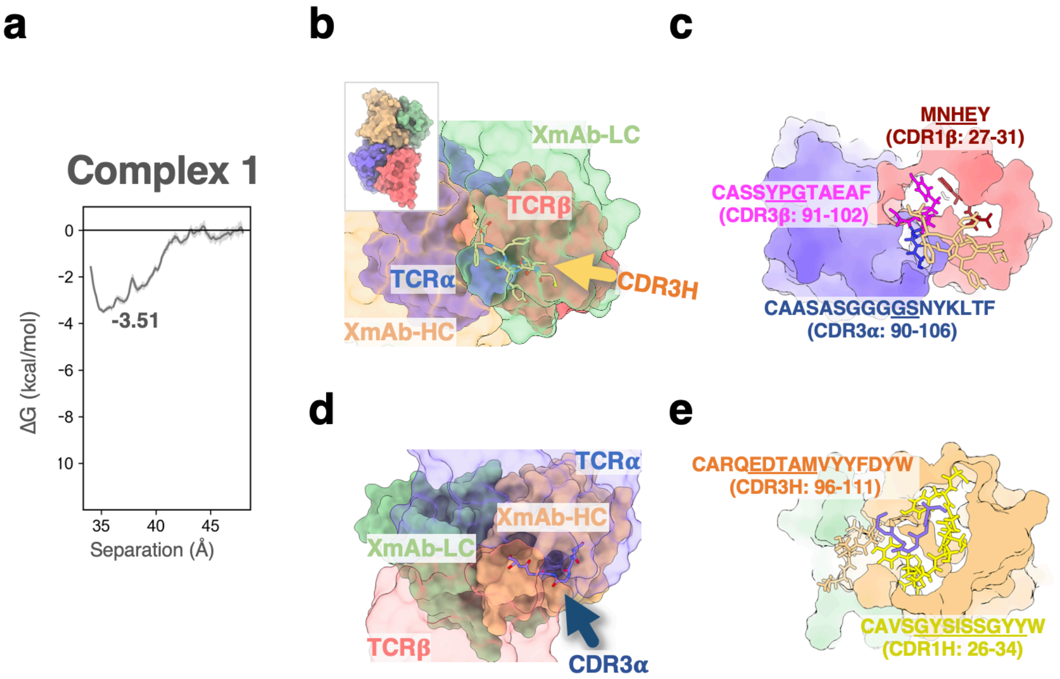


**Extended Data 9**. **Binding of the x-mAb to TCRαβ in Complex 1**. (**a**) Binding free energy for Complex 1. (**b**) Top view of the molecular surface of TCRαβ in Complex 1. The representation and coloring schemes are similar to those in Fig. 8. (**c**) CDR loops of the TCRαβ that interact with the CDR3H of the x-mAb are specifically colored and labelled. Residues that are in direct contact with the CDR3H are underlined. (**d**) Top view of the molecular surface of x-mAb in Complex 1. The representation and coloring schemes are similar to those in Fig. 8. (**e**) CDR loops of the x-mAb that interact with the CDR3α of the TCRαβ are specifically colored and labelled. Residues that are in direct contact with the CDR3α chain are underlined.

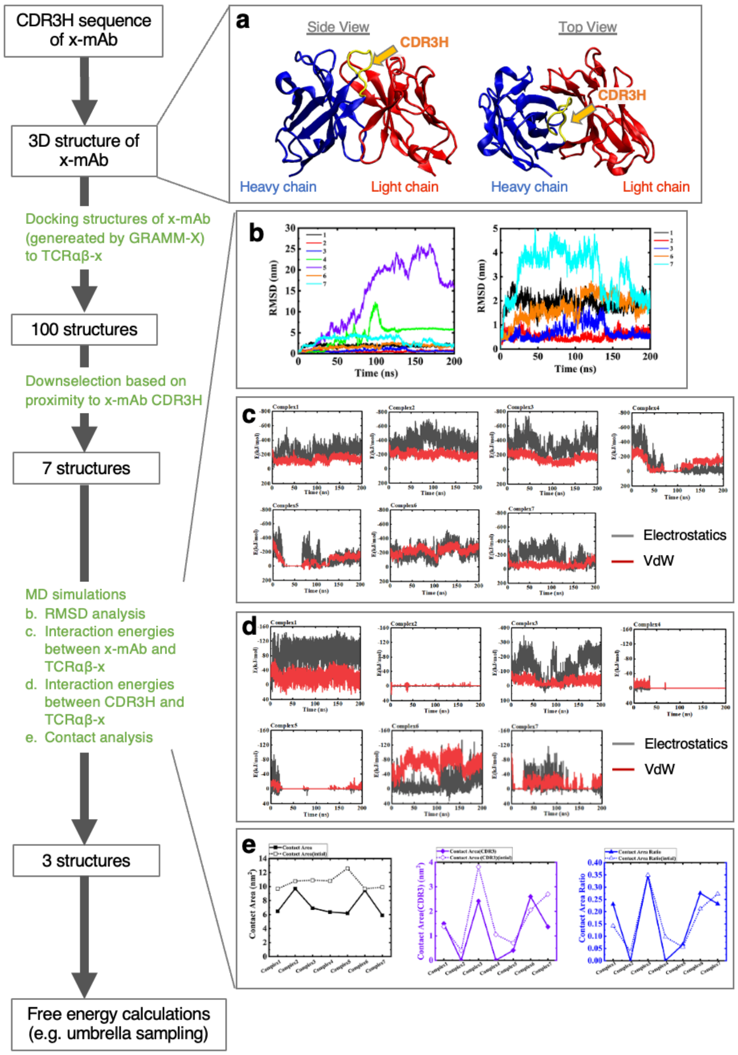


**Extended Data 10. The flowchart of TCRαβ/x-mAb complex modelling.** (**a**) The 3D structure of the x-mAb was built using previously published amino acid sequences. The light and heavy chains of the x-mAb are colored in red and blue, respectively. The CDR3 sequence (DTAMVYYFD) of the heavy chain is colored in yellow. (**b**) **Left**, RMSD analysis of MD simulations of all seven complexes. All complexes were aligned with reference to the TCRαβ. Large fluctuations correspond to Complex 4 and Complex 5. **Right**, RMSD profiles of different complexes after excluding Complex 4 and 5. (**c**) Interaction energies between the x-mAb and TCRαβ for all seven modelled complexes. Electrostatic and VdW interactions are shown in black and red, respectively. Interaction energies close to zero indicate no interaction between the two molecules. (**d**) Interaction energies between the CDR3H region of the x-mAb and TCRαβ for all seven modelled complexes during MD simulations. Electrostatic and VdW interactions are shown in black and red, respectively. Interaction energies close to zero indicate no interaction between the two molecules. (**e**) Contact area analysis. The cutoff for defining contact is 4 Å. The dashed and solid lines indicate the contact area before and after MD simulations, respectively. The contact area ratio refers to the contact area between the CDR3H region of the x-mAb and TCRαβ divided by the contact area between the x-mAb and the TCRαβ.
