## Supplemental table 13 for "Apposed networks of interacting TCRs and BCRs exhibiting mosaicked CDR3 sequences made of fixed junctional motifs"

**Supplementary Table 13: Additional examples of the CASSPGT and PGTEAFF in germline-encoded proteins**

| CASSPGT motifs in germ-line encoded proteins | | | | |
| --- | --- | --- | --- | --- |
| Organism | Common name | Gene | Position | Acc No |
| Bactrocera oleae | **Olive fruit fly** | Hemocytin | 371 to 377 | XP_014086275 |
| Cygnus color | Saker falcon bird | Fanconi anemia core complex-associated protein 100 | 398 to 404 | XM_027801266.1 |
| Penelope pileata | White crested guan (bird) | FP100 protein (found in many other species) | 196 to 202 | NXC43273 |
| Flavobacteriales | Bacteria | T9SS type A sorting domain-containing protein | 626 to 632 | JABION010000123.1 |

| CASSPGT-encoding nucleotides (tgt gcc agc agc ccc ggg atc) in germ-line proteins | | | | |
| --- | --- | --- | --- | --- |
| Organism | Common name | Gene | Position | Acc No |
| Aminobacter sp. | Bacteria | MSH1 chromosome, complete genome | 3155913 to 3155931 | CP028968 |
| Uncultured bacterium | Bacteria | Uncultured bacterium clone 16S(V3-V4)-178 16S ribosomal RNA gene | 154 to 172 | KX243731 |
| Streptomyces sp. | Bacteria | Streptomyces sp. RerS4 chromosome, complete genome | 2912290 to 2912308 | CP097322 |

| PGTEAFF-encoding nucleotides (ccc ggg act gaa gct ttc ttt) in germ-line proteins | | | |
| --- | --- | --- | --- |
| Organism | Common name | Position | Acc No |
| Apeira syringaria genome assembly, chromosome: 2 | Lilac beauty (moth) | 33020933 to 33020915 (inverted) | OW203656 |
| Agriopis aurantiaria genome assembly, chromosome: 13 | Scarce Umber (Moth) | 13156160 to 13156178 | OU611994 |
| Lithophane socia genome assembly, chromosome: 21 | the pale pinion (Moth) | 8139076 to 8139093 | OX383272 |
| ipistrellus pipistrellus genome assembly, chromosome: 15 | common pipistrelle (bat) | 6660862 to 6660880 | LR862371 |

| PGTEAFF motifs in germ-line encoded proteins | | | | |
| --- | --- | --- | --- | --- |
| Organism | Common name | Gene | Position | Acc No |
| Electrophorus electricus | Electric eel | centrosomal protein of 192 kDa isoform X1 | 1417 to 1423 | XP_035386389 |
| Scleropages formosus | Asian bony tongue | rho GTPase-activating protein 32 isoform X1 | 1365 to 1371 | XP_029107326 |
| Clostridiales bacterium | Bacteria (gut metagenome) | lycosyl hydrolase 115 family protein | 230 to 236 | MCM1448794 |
| MULTISPECIEs of bacteria | Bacteria | DEAD/DEAH box helicase [ | 135 to 141 (Pseudomonas) | WP_216705103 |
